# Region-specific reversal of epidermal planar polarity in the fancy *rosette* mouse

**DOI:** 10.1101/2023.07.27.550849

**Authors:** Maureen Cetera, Rishabh Sharan, Gabriela Hayward-Lara, Brooke Phillips, Abhishek Biswas, Madalene Halley, Evalyn Beall, Bridgett vonHoldt, Danelle Devenport

## Abstract

The planar cell polarity (PCP) pathway collectively orients thousands of cells with respect to a body axis to direct cellular behaviors that are essential for embryonic morphogenesis. Hair follicles of the murine epidermis provide a striking readout of PCP activity in their uniform alignment along the entire skin surface. Here, we characterize, from the molecular to tissue-scale, PCP establishment in the *rosette* fancy mouse, a natural variant with posterior-specific whorls in its fur, to understand how epidermal polarity is coordinated across the tissue. We find that embryonic hair follicles of *rosette* mutants emerge with reversed orientations specifically in the posterior region, creating a mirror image of epidermal polarity. The *rosette* trait is associated with a missense mutation in the core PCP gene *Fzd6*, which alters a consensus site for N-linked glycosylation and inhibits its membrane localization. Unexpectedly, this defect in Fzd6 trafficking, observed across the entire dorsal epidermis, does not interfere with the ability of other core PCP proteins to localize asymmetrically. Rather, the normally uniform axis of PCP asymmetry is disrupted and rotated in the posterior region such that polarity is reflected on either side of a transition zone. The result is a reversal of polarized cell movements that orient nascent follicles, specifically in the posterior of the embryo. Collectively, our multiscale analysis of epidermal polarity reveals PCP patterning can be regionally decoupled to produce the unique posterior whorls of the fancy *rosette* mouse.

**Summary:** Region-specific rotation of the Planar Cell Polarity axis reverses posterior hair follicles in the fancy *rosette* mouse.

## Introduction

Epidermal appendages such as hair, feathers, scales, and bristles are uniformly aligned along an animal’s body. This tissue-level pattern is a striking example of planar cell polarity (PCP) in which thousands of cells collectively orient in a common direction (Devenport and Fuchs 2008). The remarkable scale of this coordination is achieved through cell-to-cell communication mediated by a set of highly conserved membrane-associated proteins known as the core PCP pathway. Through a process that is not well understood, core PCP proteins ‘read’ long-range directional cues to orient cell polarity relative to the body axes (Fisher and Strutt 2019). The PCP machinery functions in a vast array of diverse cell types and tissues to drive collective behaviors including neural tube closure, organization of stereocilia bundles, and directional cilia beating (Butler and Wallingford 2017; Davey and Moens 2017). Therefore, deciphering mechanisms controlling PCP in highly accessible tissues like the surface ectoderm can yield fundamental insights into PCP function in other, less accessible but vital processes that shape tissues and organs during embryonic development. A hallmark of PCP organization is its long-range coordination with a body or tissue axis. Despite this common feature, the mechanisms that link the axis of PCP asymmetry to a body axis remain elusive.

The core components of the PCP pathway were discovered using forward mutagenesis screens in *Drosophila* where misoriented wing hairs and cuticular bristles served as phenotypic indicators of defective planar polarity. The PCP genes *frizzled*, *disheveled*, *prickle*, *starry night* (aka *flamingo*), and *van gogh* were aptly named for their mutant phenotypes of disordered or swirled hair and bristle patterns (Wolff and Rubin 1998; Taylor et al. 1998; Gubb and García-Bellido 1982; Krasnow, Wong, and Adler 1995). In other species where mutagenesis is less practical, natural genetic variation can provide a powerful means to uncover new genes or novel mutations in developmental pathways in an unbiased way. For example, animal breeders have selected for spontaneous mutations in birds and rodents that give rise to beautiful feather and hair patterns such as the collar of the crested pigeon or the whorls of the Abyssinian guinea pig (Shapiro et al. 2013; Castle 1905). In both cases, the polarity of epidermal appendages no longer uniformly aligns with the anterior-posterior (AP) body axis, suggesting tissue-level planar polarity is disrupted. PCP is an ideal developmental pathway to genetically dissect using natural variation because misaligned epidermal structures are easily visualized by the naked eye, producing traits that are desirable to breeders. Identifying the sequence variants associated with unique epidermal patterns will likely uncover novel mutations in the PCP pathway itself or new genes required for PCP patterning.

The “*rosette*” trait in the mouse, *Mus musculus,* is a unique coat pattern where the dorsal fur forms two large symmetrical whorls, or rosettes, in the posterior half of the body (Figure 1A). The phenotype appeared in a British fancier’s colony in the 1960s and is now a coveted trait in the breeding community (King 1975; Wallace 1971). The recessive phenotype can be combined with a number of different coat colors, textures, and lengths to create truly unique animals. The *rosette* trait shares features with the hair phenotypes caused by mutations in PCP pathway genes in laboratory mice, but the region-specific nature of the coat pattern sets it apart from previously described PCP mutants. Of the PCP mutant strains that survive to adulthood, hair whorls typically arise over the entire skin surface (Ravni et al. 2009; Guo, Hawkins, and Nathans 2004). Further, loss-of-function mutations in PCP genes tend to cause neural tube defects and embryonic lethality requiring skin-specific knockouts to observe hair patterning defects (Cetera et al. 2017; Curtin et al. 2003; Murdoch et al. 2001; Kibar et al. 2001). These key differences suggest *rosette* is a novel variant that alters PCP signaling in a region-specific manner.

**Figure 1.**
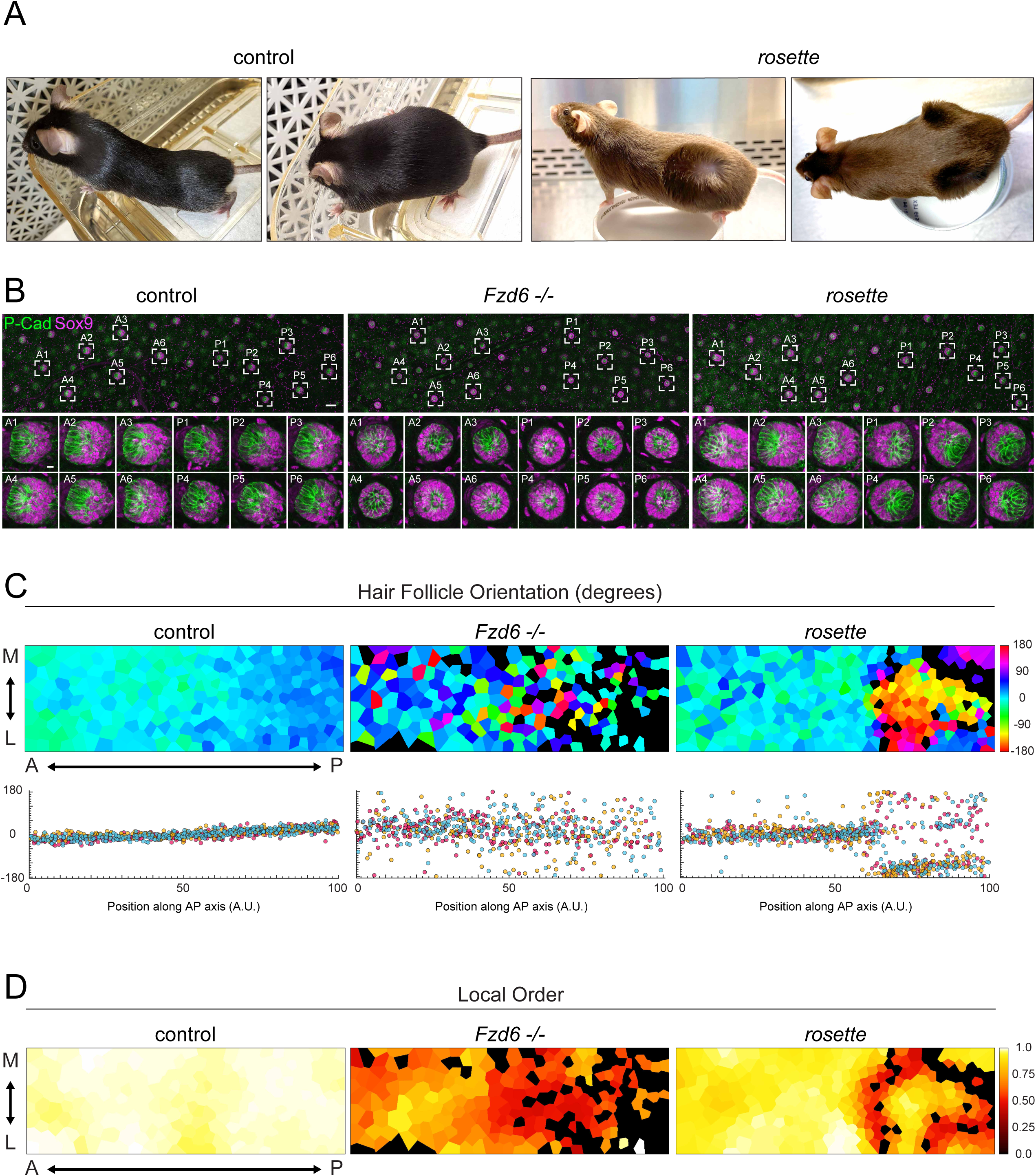
Posterior hair follicles emerge with reversed orientations in *rosette* mutants. (A) Hair patterns in adult stage C57BL/6 controls (left) and the fancy *rosette* mouse (right). Two large symmetric hair whorls are present specifically in the posterior region of *rosette* animals. (B-D) Orientation of emerging hair follicles in control (left), *Fzd6-/-* (center), and *rosette* (right) embryonic skins at E15.5. (B) Representative scanning confocal projections of whole mount, dorsal skins labeled with asymmetrically distributed follicle markers, P-Cad (green) and Sox9 (magenta). Scale bar, 100 microns. Zoomed views of individual follicles are shown below. “A1, A2, etc.” indicates anterior follicles and “P1, P2, etc.” indicates posterior follicles from the corresponding image above. Scale bar, 10 microns. (C, top) Voronoi diagram representing the orientation of follicles across an entire flank of skin by color (cool colors= anteriorly oriented follicles, warm colors= posteriorly oriented follicles, black=unpolarized). X-axis, anterior-posterior. Y-axis, medial-lateral. (C, bottom) Graph showing the orientation of individual follicles with respect to their AP position. Three different colors indicate three embryos per genotype. (D) Voronoi diagram representing the local order of follicle orientations in a radius of 3 follicles as described in the Methods section (high coordination=white, low coordination= red, unpolarized follicles=black).

Here, we describe how the extraordinary *rosette* coat pattern emerges during skin development at the cellular, appendage, and tissue-wide scales, and show that the whorls are the result of aberrant PCP establishment. We find posterior whorls develop from hair follicles that emerge with reversed orientations during embryonic stages, the time at which the PCP pathway initially acts to polarize nascent hair follicles. Genetic mapping revealed that mice displaying the *rosette* trait are homozygous for a novel missense mutation in the *Fzd6* locus, a known core component of the PCP pathway. We show that the missense mutation, which is predicted to inhibit N-linked glycosylation of the Fzd6 extracellular domain, interferes with proper Fzd6 trafficking through the secretory system and prevents its delivery to the plasma membrane. Under this condition, Fzd6 accumulates intracellularly in the skin epithelium while the other core PCP components become polarized but are not uniformly coordinated across the tissue. In the posterior, the PCP axis rotates a full 180 degrees, reversing the orientation of collective cell movements that polarize the hair follicle. Collectively, our analysis of epidermal polarity in *rosette* embryos reveals Fzd6 plays a role in directing the orientation of PCP in the posterior region of the skin and suggests that different skin regions can independently establish an axis of PCP asymmetry.

## Results

### Posterior hair follicles emerge with reversed orientations in *rosette* mutants

The “*rosette*” fancy mouse is prized for its distinctive coat marked by posterior-specific whorls (Figure 1A). Despite gaining popularity among fanciers, *rosette* mice were not available to the scientific community. This led us to attend regional fancy mouse shows where we networked with the breeding community to obtain and rederive the whorled mice in a laboratory setting. Although swirling hair patterns in mice are indicative of a defect in the PCP pathway, the regionality of the *rosette* phenotype is highly unusual as PCP is required to align hair follicles across the entire skin (Ravni et al. 2009; Cetera et al. 2017; Guo, Hawkins, and Nathans 2004). Hair follicle polarity is established during embryonic development shortly after hair placodes are specified from the basal progenitor layer of the embryonic epidermis (Devenport and Fuchs 2008). Asymmetrically localized PCP proteins direct polarized cell movements within each placode, causing the multicellular structure to tilt and grow in an anterior orientation (Cetera et al. 2018). In the absence of PCP function, hair placodes emerge with randomized orientations or they fail to polarize altogether and grow vertically into the dermis (Devenport and Fuchs 2008; Cetera et al. 2017; Wang, Chang, and Nathans 2010; Dong et al. 2018; Basta et al. 2023). To determine whether the *rosette* phenotype arises during the initial stages of hair follicle polarization, we quantified hair placode orientations in *rosette* mutant embryos at E15.5 and compared these to wild type controls as well as a loss-of-function PCP mutant (*Fzd6-/-*).

The orientation of embryonic follicles can be determined by examining the relative positions of P-Cadherin (P-Cad) and Sox9-expressing cells within individual follicles. In wild type embryos, these two cell populations are asymmetrically positioned at the anterior and posterior sides of each follicle with P-Cad expressing cells oriented toward the head of the animal (Figure 1B, A1-6, P1-6, Figure S1A). By contrast, in *Fz6-/-* mutant follicles, P-Cad and Sox9 expressing cells are either mispositioned relative to the AP axis (A1, A2, A3, P6) or unpolarized (A4, A5, P1, P2, P3, P4, P5) with Sox9 cells encircling a central cluster of P-Cad cells (Cetera et al. 2017; Wang, Chang, and Nathans 2010). The embryonic hair pattern of *rosette* mutants, by contrast, is distinct. Follicles located in the anterior half of the embryo position P-Cad expressing cells anteriorly (A1-6) as they do in wild type controls, while follicles in the posterior region of the embryo are reversed, with P-Cad and Sox9 expressing populations inverted relative to the AP axis (P1-6).

To characterize tissue-scale hair follicle patterns in each genotype, we used a semi-automated method to measure follicle orientations based on P-Cad and Sox9 distributions (Basta et al. 2023). We displayed this information at the tissue level in Voronoi diagrams where each polygon represents a follicle that is color-coded according to its orientation from 0 to 360 degrees (Figure 1C, top; cool colors = anteriorly oriented; warm colors = posteriorly oriented; black= unpolarized, Figure S1A). To determine reproducibility between individuals, follicle orientations were plotted relative to their AP positions across multiple embryos (Figure 1C, bottom). In control embryos, follicles point anteriorly and are well aligned between +45 and -45 degrees. Follicle orientations in *Fzd6-/-* mutants are much more broadly distributed, ranging from +180 to -180 degrees, with unpolarized follicles scattered across the tissue. In *rosette* embryos, the majority of follicles correctly align between +45 and -45 degrees in the anterior half of the skin, while follicles in the posterior region are reversed, predominantly aligning near +180 and -180 degrees.

As a third metric to describe the collectivity of follicle orientations, we defined an order parameter that compares the relative angles of neighboring follicles (Figure 1D, Equation 1 in Materials and Methods). Parallel hair follicles are highly ordered with an order parameter of one (white) while antiparallel hair follicles have an order parameter of zero (dark red). In control embryos, local order is high across the entire skin (white, yellow) whereas in *Fzd6-/-* mutants, local order is low, with each follicle appearing to orient independently of its neighbors (orange, red). By contrast, local order is high in both the anterior and posterior regions of *rosette* embryos (yellow) with the exception of a sharp zone of low order surrounding the reversed patch of posterior follicles. In this region, follicles are changing their orientation or growing straight down (black) resulting in the low coordination between neighboring follicles (red). Together, these data demonstrate that hair follicle polarity is altered in a region-specific manner in *rosette* mutants, and these defects arise during the earliest stages of hair follicle polarization. Moreover, the reversed orientations and high local order of follicles in *rosette* mutants comprise a phenotype that is distinct from a loss-of-function PCP mutant.

### Coordinated hair follicle reversal persists into postnatal stages

Misoriented hair follicles in PCP mutants are not static but instead have the remarkable ability to rotate up to 180 degrees. These rotations minimize the angular difference between adjacent hair follicles during postnatal stages, which ultimately gives rise to the elaborate whorls, ridges, and crosses that mark the coats of certain PCP mutants (Wang, Chang, and Nathans 2010; Wang, Badea, and Nathans 2006; Cetera et al. 2017). Postnatal follicle rotations can also correct an initially disordered follicle pattern. For example, in *Fzd6-/-* mutants, initially misoriented hairs rotate and correct their alignment along the AP axis and by postnatal day 7 (P7), their coats are indistinguishable from wild type animals mutants (Wang, Chang, and Nathans 2010; Wang, Badea, and Nathans 2006; Cetera et al. 2017). To examine how the *rosette* hair pattern evolves and refines during postnatal stages, we measured hair follicle angles and local order across the entire dorsal skin at postnatal day 4 (P4) when *Fzd6-/-* follicles are in the process of reorienting. At this stage in *Fzd6-/-* mutants, most follicles have rotated to face anteriorly or point slightly away from the midline, particularly in the posterior region, as previously described (Figures 2A, S2, cool colors, Figure S1B)(Cetera et al. 2017). Furthermore, previously uncoordinated follicles at embryonic stages are highly ordered by P4 (Figures 2B, S2, white/yellow). In contrast, follicles in the posterior region of *rosette* mutants retained their reversed, posterior-facing orientations even at P4, when follicles have the ability to rotate into correct orientations (Figures 2A, S2, warm colors). The zones of high and low local order that were observed in embryonic stages were also maintained at P4. Interestingly, all previously reported PCP mutants display rear paw whorls that are maintained throughout postnatal stages, but they are not present in *rosette* animals (Figure 2C) (Cetera et al. 2017; Guo, Hawkins, and Nathans 2004; Basta et al. 2023; Ravni et al. 2009). These data show misoriented follicles in the *rosette* mutant fail to correct their orientation during early postnatal stages and *rosette* mutants lack the characteristic PCP-mutant paw whorls.

**Figure 2.**
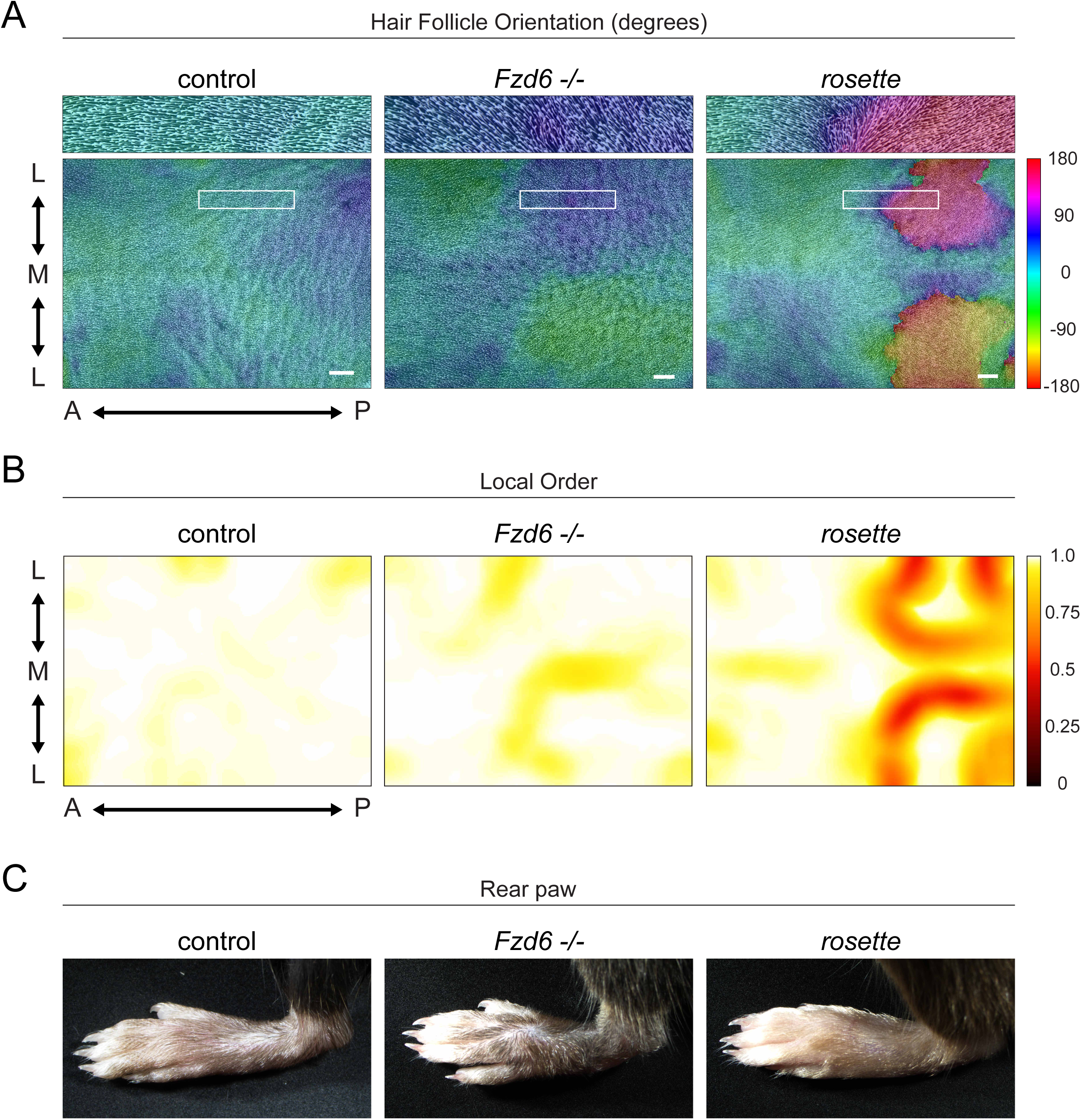
Coordinated hair follicle reversal persists into postnatal stages. (A) Representative images of cleared, flat-mounted dorsal skins at postnatal day 4 from *rosette/+* controls (left), *Fzd6-/-* (center), and *rosette* (right). Hair follicle orientation is indicated by the color overlaid on the hair follicle image (cool colors= anteriorly oriented follicles, warm colors= posteriorly oriented follicles). Scale bar= 1mm. (B) Local order maps corresponding to the same images in A. Local order is calculated across a radius of 200 pixels containing approximately 200-300 follicles (high order= white, low order= red). AP, anterior-posterior. ML, medial-lateral with M indicating the midline. (C) Rear paw whorls from indicated genotypes showing whorls are present in Fzd6-/- but absent in controls and *rosette* mutants.

### The *rosette* phenotype correlates with a missense mutation in Fzd6

To uncover genetic elements associated with the *rosette* phenotype, we leveraged the diversity between the genetic backgrounds of fancy and laboratory mice. Although common lab strains originated from fancy mice, they are derived from a limited number of founders and are highly inbred (Phifer-Rixey and Nachman 2015). The background of fancy mice, however, is mixed and animals are derived from multiple sources to obtain the unique combination of traits desired by breeders. The phenotypic animals in our colony are descendants of a single *rosette* founder crossed to the C57BL/6 inbred lab strain. Using a Mouse Diversity Genotyping Array, we identified SNPs in the fancy background that co-segregate with the appearance of posterior whorls (H. Yang et al. 2009). We analyzed 91 animals in total including the founder, its *rosette* sibling, two C57BL/6 animals and descendants from the fancy founder with increasing levels of C57BL/6 ancestry (Tables S1, S2). After applying a minor allele frequency of <0.3%, we retained 67,589 SNP loci that capture the gradient of C57BL/6 ancestry in Principle Component Analysis 1 showing the maximum variance is dependent on the genetic background (Figure 3A). We used a linear mixed model to identify SNP variants associated with the whorl phenotype and used the likelihood ratio test (LRT) for significance testing (X. Zhou and Stephens 2012). We prioritized SNPs with allelic effect (ß) values in the lower or upper 5th-percentile of the distribution and considered these putative outlier SNPs. By integrating the LRT p-value and ß values, we identified a candidate region on chromosome 15 that contained 132 genes, including *Fzd6* (Figure 3B). Despite the distinct hair patterning defects displayed in *Fzd6-/-* and *rosette* mutants, we reasoned that a novel mutation in *Fzd6* might contribute to the unique *rosette* phenotype.

**Figure 3.**
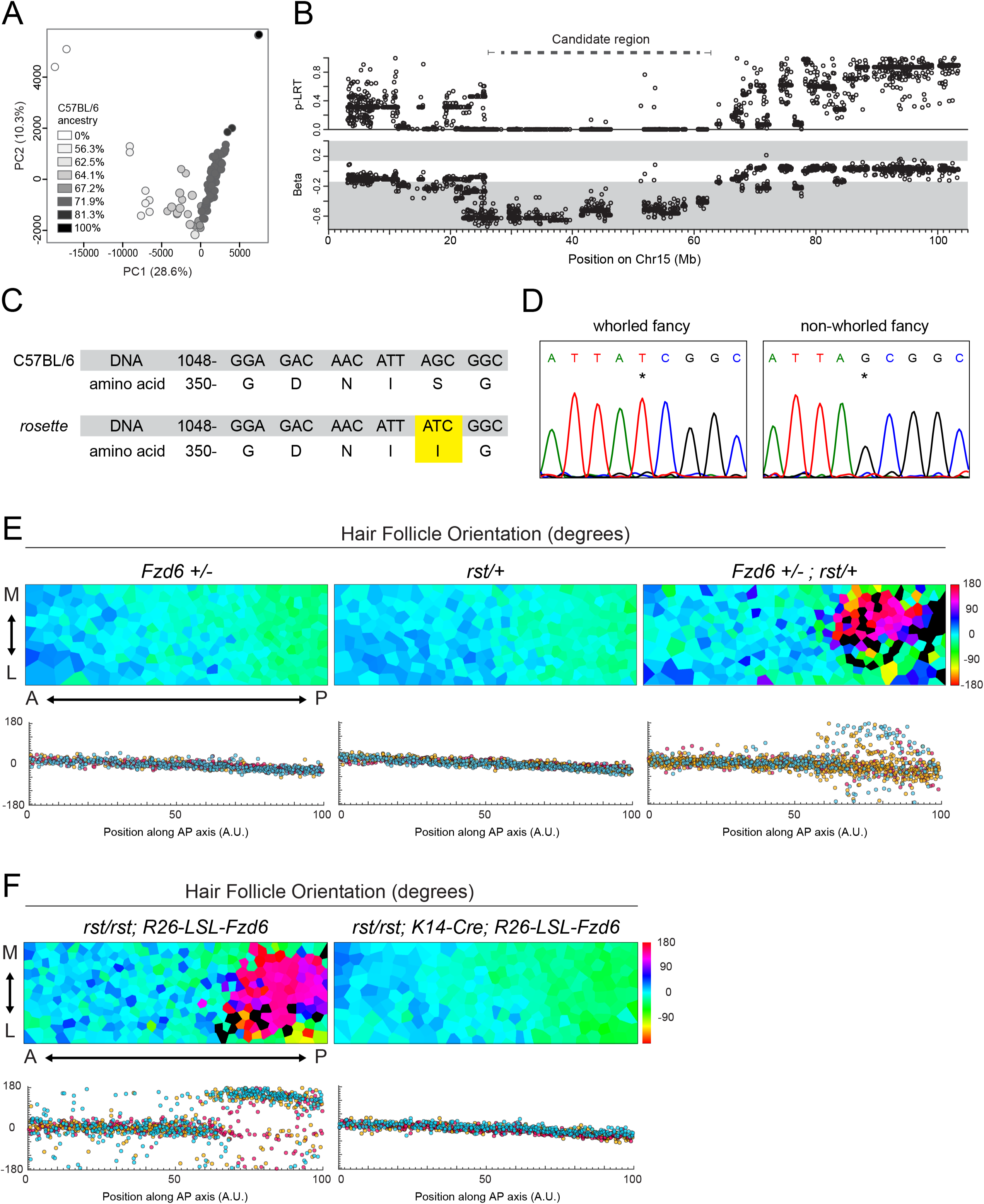
The *rosette* phenotype correlates with a missense mutation in *Fzd6*. (A) PCA of the filtered SNP data set for all mice analyzed. Circle shading indicates the C57BL/6 ancestry for each animal. Percent variation for the first two axes are included. (B) Candidate region associated with the whorled phenotype on chromosome 15 (dashed line) identified by outlier loci from the likelihood ratio test statistic (top panel) and outlier ß values (lower panel, gray shading). (C) A novel missense mutation in the *rosette* mutant compared to the C57BL/6 reference sequence. G1061>A results in an amino acid change in the Fzd6 protein, S354I (yellow). (D) Sanger sequencing data showing *rosette* animals from outside of our colony have the same point mutation (left, asterisk) while non-whorled fancy mice do not (right, asterisk). (E) *rosette* genetically interacts with *Fzd6*. (E, top) Voronoi diagrams representing embryonic hair follicle orientations across entire skin flanks of heterozygous *Fz6 +/-* (left), *rosette/+* (*rst*, center), or *Fzd6+/-; rst/+* (right) embryos at E15.5 (cool colors= anteriorly oriented follicles, warm colors= posteriorly oriented follicles, black=unpolarized). AP, anterior-posterior. ML, medial-lateral. (E, bottom) Graph showing the orientation of individual follicles with respect to their AP position in embryos of the indicated genotypes. Three different colors indicate three embryos per genotype. (F) Rescue of the *rosette* phenotype upon epidermal expression of wild-type *Fzd6*. (F, top) Voronoi diagrams representing embryonic hair follicle orientations across entire skin flanks of homozygous *rst/rst* embryos with (right) or without (left) expression of wild-type *Fzd6* driven by *K14-Cre*. (cool colors= anteriorly oriented follicles, warm colors= posteriorly oriented follicles, black=unpolarized). AP, anterior-posterior. ML, medial-lateral. (F, bottom) Graph showing the orientation of individual follicles with respect to their AP position in embryos of the indicated genotypes. Three different colors indicate three embryos per genotype.

Exon sequencing of the *Fzd6* gene in phenotypic *rosette* animals identified four point mutations not present in the C57BL/6 reference genome, three of which are known SNP variants found between lab strains (Table S3)(Lilue et al. 2018). The fourth is a novel missense mutation, G1061>A, that results in a serine to isoleucine substitution at amino acid 354 (S354I, Figure 3C). To determine if the missense mutation was linked to the *rosette* phenotype in mice outside of our colony, we sequenced the same region in three unrelated whorled animals obtained from an independent breeder. All three animals had the G1061>A mutation in *Fzd6* (Figure 3D). To rule out the possibility that G1061>A arose in the background of fancy animals independent from their whorled status, three non-whorled animals from the breeder’s colony were sequenced. None carried the mutation revealing the G1061>A mutation in Fzd6 correlates with the *rosette* phenotype (Figure 3D). To test for a genetic interaction between *rosette* and *Fzd6*, phenotypic *rosette* animals were crossed to *Fzd6-/-* homozygotes and their heterozygous progeny were scored for hair polarity defects. If the S354I substitution impairs Fzd6 function, it should fail to complement the *Fzd6* null allele (Figure 3E, Figure S1A). We found that while hair patterning is normal in embryos heterozygous for *Fzd6+/-* or *rosette/+* alone, in combination, follicles in the posterior region are misoriented indicating the S354I mutation likely alters Fzd6 function.

Interestingly, the S354I allele shows a stronger phenotype in homozygous animals than when heterozygous with the null allele (Figure 1C compared to 3E). If the S354I substitution is a hypomorph, we would expect a stronger, not weaker, phenotype in combination with the null allele. One potential explanation is that a recessive modifier is present in the *rosette* background that is missing in our *Fz6* knockout lab strain. A previous study showed that the *Fzd6-/-* follicle pattern can be enhanced when combined with a naturally occurring deletion of exon 5 in *Astrotactin2* found in the129X1/SvJ lab strain (H. Chang et al. 2015). However, we did not detect the deletion in the fancy *rosette* background (Figure S3). Although this result does not rule out that another modifier could contribute to the phenotype, it does show reversal cannot be explained by combining previously described alleles. If impaired Fzd6 function underlies the coat patterning phenotype of *rosette* mutants, expressing an exogenous copy of wild-type *Fzd6* should rescue the hair follicle reversals. We therefore overexpressed Fzd6 in *rosette* mutants by driving expression of the *Rosa26 lox-stop-lox Fzd6WT* transgene with a skin-specific driver, *K14-Cre* (Hua et al. 2014; Vasioukhin et al. 1999) Figure 3F, Figure S1A). In the absence of *K14-Cre*, the polarity of *rosette* mutant hair follicles was unchanged; follicles in the posterior region were reversed. However, upon expression of wild-type *Fzd6* with *K14-Cre*, all follicles were collectively oriented toward the anterior of the animal, restoring the wild-type pattern. Together, data from our mapping, genetic interaction, and rescue experiments strongly suggest that altered Fzd6 function contributes to the *rosette* phenotype.

### Membrane localization of Fz6 is lost in *rosette* mutants

Membrane proteins require N-linked glycosylation for proper protein folding and export from the ER. Fzd6 has two predicted N-linked glycosylation sites at luminal positions N38 and N352. When both sites are mutated to alanine and expressed in Cos7 or HEK293 cells, Fzd6^N38A;N352A^ is retained in the ER (Tang et al. 2020). The S354I mutation identified in *rosette* mutants alters the consensus sequence N-X-S/T which is essential for N-linked glycosylation. The consensus sequence is found in extracellular loop two and although it is not present in *Drosophila*, it is conserved among tetrapod species suggesting functional importance (Figure S4A). If glycosylation at N352 is inhibited in *rosette* mutants, we might expect to see a change in Fzd6 localization. Fzd6 normally localizes to the plasma membrane of basal epidermal cells where it colocalizes with E-cadherin at intercellular junctions (Figure 4A, top, Figure S1A). As expected for a defect in ER export, colocalization between Fzd6 and E-Cadherin is lost in *rosette* mutants and instead, Fzd6 localizes intracellularly to the space between the nucleus and the plasma membrane (Figure 4A, bottom, Figure 4B). Interestingly, although hair follicle polarity reversal only occurs in the posterxior region of *rosette* embryos, we found that Fzd6 is mislocalized across the entire epidermis (Figure 4B).

**Figure 4.**
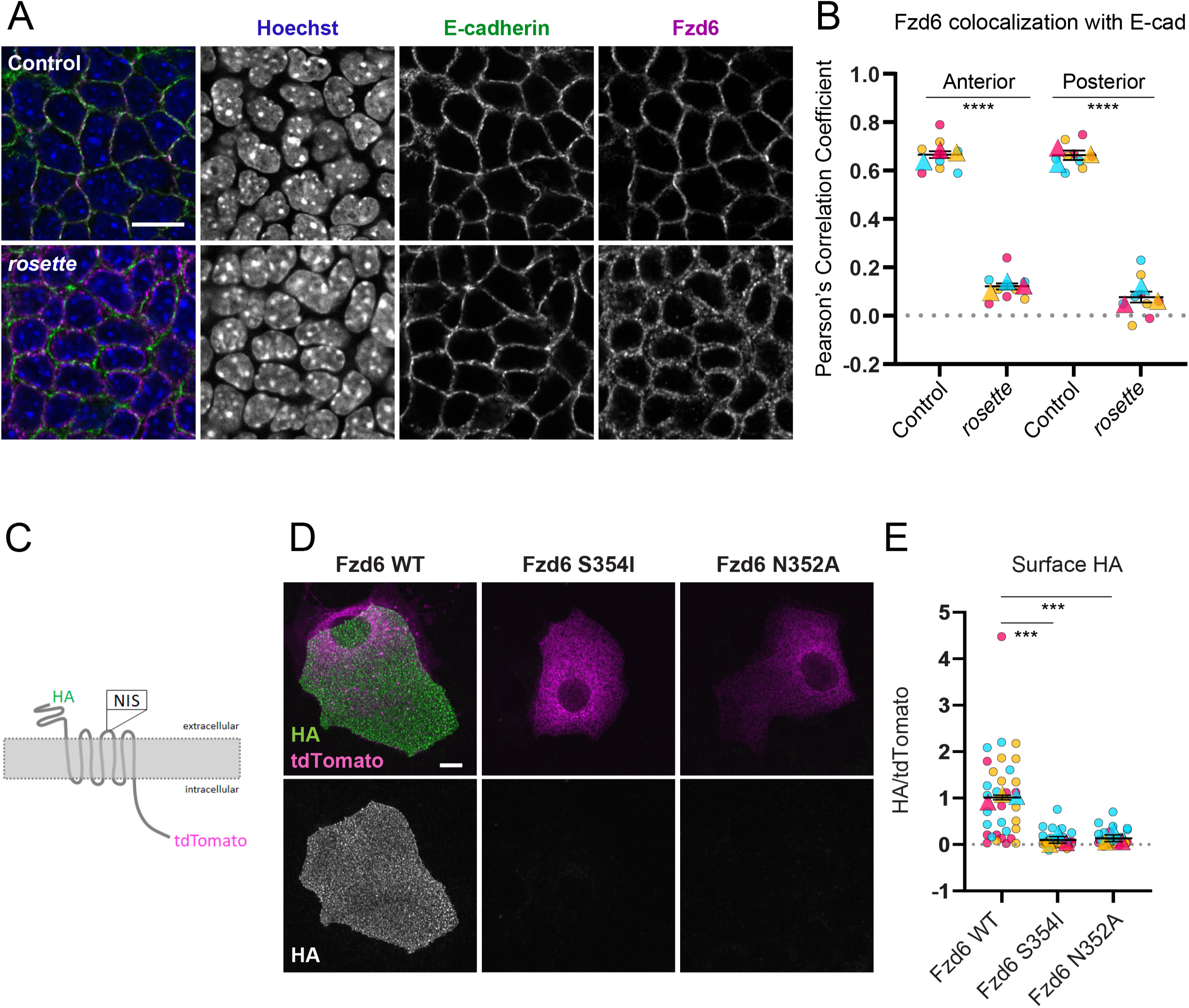
Membrane localization of Fz6 is lost in *rosette* mutants. (A) Scanning confocal images of whole mount epidermis labeled with E-cadherin and Fzd6 antibodies and Hoechst in control (top) and *rosette* (bottom) embryos at E15.5. Single optical slices through the basal layer are shown. Scale bar= 10 microns (B) Colocalization of E-cadherin and Fzd6 in non-nuclear areas measured by Coloc2 in ImageJ and represented as Pearson’s correlation coefficient. Three 30×30 micron images were analyzed in the anterior and posterior regions of each skin. Three animals were analyzed per genotype, represented by unique colors. Circles indicate a single analyzed image and the triangle is the average/ animal. Mean and SEM across all animals per group are indicated. Two-tailed t-test: p< 0.00001, anterior, p<0.00005, posterior. (C) Schematic of Fzd6 constructs tagged with extracellular HA and intracellular td-Tomato used for keratinocyte transfection and surface labeling. The NIS sequence is altered in mutants as indicated in D. (D) Keratinocytes transfected with HA-Fzd6-tdTomato constructs (WT, S354I, N352A) and labeled with anti-HA antibodies without permeabilization. Maximum projections of widefield stacks are shown. Scale bar, 10 microns. (E) Quantification of surface HA levels relative to the tdTomato signal. WT n=36, N354I n=34, N352A n=35 cells from 3 experiments. Each experiment is shown in a different color. Replicates are circles and triangles are the average of the experiment. Mean and SEM are shown. Two-tailed t-test: p=0.00059 (WT vs S354I) p=0.00099 (WT vs N352A).

To determine if the S354I mutation itself is responsible for Fzd6 intracellular accumulation as opposed to another genetic element in the *rosette* background we introduced the mutation into a Fzd6 construct with intracellular tdTomato and extracellular HA tags and transfected control or mutant Fzd6 constructs into cultured mouse keratinocytes (Figure 4C). By performing anti-HA staining without membrane permeabilization, we could selectively label the surface exposed Fzd6 extracellular domain and compare the levels to total Fzd6-tdTomato. In keratinocytes expressing wild-type HA-Fzd6-tdTomato, HA labeling was abundant on the cell surface (Figures 4D, E). By contrast, HA labeling was not detectable on the surface of cells expressing HA-Fzd6-S354I-tdTomato and the tdTomato signal was localized to an intracellular tubular network that likely corresponds to the ER. As a control, we performed anti-HA labeling in the presence of detergent to ensure the mutant constructs functioned as expected and found that HA localized intracellularly together with tdTomato (Figure S4B).

Next, we tested whether inhibiting glycosylation at position N352 would mimic the trafficking defect caused by the S354I mutation. We substituted asparagine 352 with alanine and performed surface HA labeling on keratinocytes expressing HA-Fzd6-N352A-tdTomato. Again, surface HA levels were not detectable and td-Tomato was localized to intracellular membranes mirroring the S354I mutation (Figures 4D, E). These results suggest the S354I mutant likely inhibits N-linked glycosylation and that blocking the post-translational modification is sufficient to prevent the delivery of Fzd6 to the cell surface.

### The axis of PCP asymmetry rotates in the *rosette* epidermis

In the wild-type epidermis, PCP proteins are polarized along AP junctions with Celsr1 displaying axial asymmetry, forming homodimers between neighboring cells, and Vangl2 and Fzd6 displaying vectorial asymmetry, polarizing toward anterior or posterior sides of the cell, respectively (Figure 5A)(Devenport and Fuchs 2008). Loss of a single PCP protein leads to uniform distribution of the remaining proteins (Devenport and Fuchs 2008; Cetera et al. 2017; Stahley et al. 2021). To understand how mislocalized Fzd6 could lead to region-specific follicle reversal in *rosette* embryos we examined the axis of Celsr1 polarity, focusing on the transition zone, where follicles switch from anterior to posterior orientations. We used QuantifyPolarity to measure the orientation and magnitude of Celsr1 polarity based on Principal Component Analysis (Figures 5B, Supplemental Figure 5)(Tan et al. 2021). In wild-type embryos, Celsr1 is enriched along AP junctions (predominantly within 30 degrees of 0), depleted from mediolateral (ML) junctions (within 30 degrees of ±90) and the magnitude of asymmetry is high. By contrast, Celsr1 is weakly polarized in no particular direction in *Fzd6-/-* mutants. In *rosette* mutants, Celsr1 is predominantly polarized along AP junctions but not as strongly as in wild-type controls.

**Figure 5.**
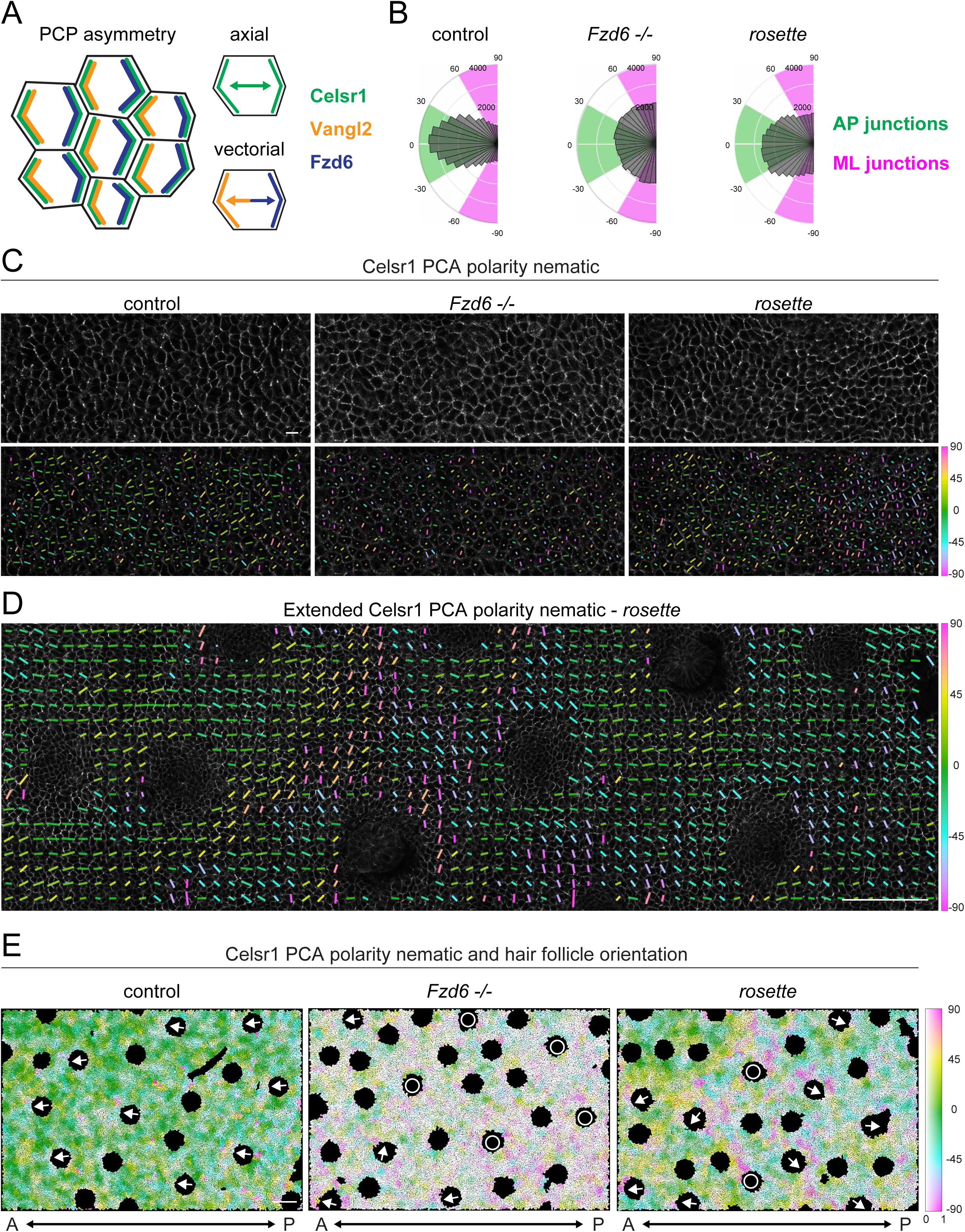
The axis of PCP asymmetry rotates in the *rosette* epidermis. (A) Schematic of PCP asymmetry in wild type basal epidermal cells. Celsr1 (green) is polarized to both anterior and posterior sides of the cell (axial) while Vangl2 (orange, anterior) and Fzd6 (blue, posterior) are polarized toward opposite sides of the cell (vectorial). (B-E) Celsr1 polarity in the basal layer of E15.5 epidermis. The areas shown are in the transition zone, near where the follicles reverse in *rosette* mutants and the corresponding area in *Fzd6-/-* and controls. (B) Circular histograms of Celsr1 polarity angles weighted by magnitude. Control n=46,256 cells, *Fzd6-/-* n=45,854 cells, and *rosette* n=48,031 cells from 3 embryos/genotype. (C, top) Scanning confocal images of whole mount epidermis labeled with Celsr1 antibodies from control (left), *Fzd6-/-* (center), and *rosette* (right) embryos. Scale bar = 10 microns. (C, bottom) Celsr1 polarity nematic calculated by PCA with QuantifyPolarity. Lines point to the sides of the cell with the highest levels of Celsr1 and are color coded by angle (green=AP, magenta=ML). Magnitude is indicated by the length of the line with longer lines indicating higher magnitudes. (D) Extended AP region showing the average polarity nematic for a zone of approximately 10 cells. Blank spaces are newly formed follicles. Scale bar =100 microns. (E) Spatial map of local Celsr1 polarity angle (color) and magnitude (saturation). Each cell is color-coded to represent the average angle and magnitude of a local area with a radius of 40 pixels (center cells and approximately 2 radii of neighbors). Black circular regions correspond to emerging hair follicles and arrows depict their orientations. White circles mark unpolarized follicles that point straight down. Blank spaces represent newly formed follicles that are too early to categorize or areas that could not be segmented. Scale bar =100 microns.

To understand how Celsr1 is organized to produce this intermediate phenotype, we used QuantifyPolarity to display the polarity nematic over each cell as a line that points to the sides of the cell with the highest level of Celsr1 with the length of the line indicating the magnitude (Figure 5C, Figure S1A). Lines were color-coded to indicate orientation (green=AP, magenta=ML). In wild-type embryos, Celsr1 is strongly polarized along AP junctions (long, green/yellow lines), whereas in *Fzd6-/-* embryos, Celsr1 is weakly polarized and randomly oriented (shorter, uncoordinated lines). Interestingly, Celsr1 is strongly polarized and locally coordinated in *rosette* mutants, but the axis of polarity is not fixed. Instead, Celsr1 rotates from mainly AP junctions (long green/yellow lines) to ML junctions (long magenta lines) forming a swirling pattern, which has never been previously described in the skin. To see if ML enrichment resolves posteriorly, we analyzed a larger region and displayed the average polarity nematic over approximately 10 cells (Figure 5D, Figure S1A). Celsr1 rotates from AP to ML and back to AP junctions (Figure 5D, green, magenta, green). To determine if the changes in the PCP axis correlate with the overall hair follicle pattern, we extended the imaging area further to encompass over 15,000 cells and 15-25 hair follicles and generated spatial maps to represent the average axis of polarity by color and magnitude by saturation (Figure 5E, low saturation=low magnitude, high saturation=high magnitude, Figure S1A). In wild-type embryos, follicles are anteriorly oriented (arrows) and Celsr1 is strongly polarized along AP junctions whereas follicles are misaligned or unpolarized (circles) in *Fzd6-/-* mutants and Celsr1 is weakly polarized in random orientations. In *rosette* mutants, Celsr1 asymmetry is locally aligned with high magnitudes which can be observed as highly saturated patches of cells of the same color. Although Celsr1 is polarized in many different orientations, it is often aligned along the ML axis (magenta) where follicles transition from anterior to posterior orientations. Further away from this zone, in both anterior and posterior directions, Celsr1 is predominantly polarized along AP junctions (0±40 degrees, green-blue, green-yellow). These data suggest that in *rosette* mutants, the axis of PCP asymmetry rotates in the transition zone and is likely reversed in the posterior by a full 180 degrees.

### PCP-directed collective cell movements are reversed in *rosette* mutants

The polarized morphogenesis of the hair placode results from PCP-directed collective cell movements (Cetera et al. 2018). If the vector of PCP asymmetry in individual cells is reversed in the posterior region of *rosette* animals such that the PCP proteins that are normally present at the anterior side of a junction are now present at the posterior side, the initial PCP-directed cell movements that establish the hair follicle tilt should also be reversed. To test this, we performed live imaging of skin explants from *rosette* mutant embryos and heterozygous controls focusing on newly formed placodes located in both anterior and posterior regions. Epithelial cells were visualized with the epidermis-specific nuclear marker, K14-H2B-GFP and/or membrane-tdTomato (Muzumdar et al. 2007; Tumbar et al. 2004). Cells were tracked over approximately 17 hours as initially radially symmetric placodes underwent polarized cell rearrangements. In control explants, cells of the placode epithelium rearrange in a counter-rotational pattern, where posterior cells converge and migrate toward the anterior (blue cells) and anterior positioned cells move outward and posteriorly (red, orange cells), as we previously described (Figures S6A, B, Supplementary Videos 1, 2, n=4 anterior, n=3 posterior, Figure S1A)(Cetera et al. 2018). In the anterior region of *rosette* mutants, cell rearrangements within placodes were similar to wild type controls (Figure 6A, Supplementary Videos 3, 4), with the exception of one placode, where cells failed to move (n=7). In the posterior region, by contrast, cell rearrangements were reversed; anterior cells converged and moved toward the posterior (red, orange cells), while posterior cells were swept outward and toward the anterior (blue cells, Figure 6B, Supplementary Videos 5, 6 n=7/9). In two placodes located in the posterior region of *rosette* mutant explants, cell movements were directed in the correct, anterior-biased orientation, likely because they were located near the zone of reversal (n=2/9). Thus, hair follicle reversal occurs due to reversed collective cell rearrangements at the initial stages of hair follicle morphogenesis, strengthening the evidence that the *rosette* mutation causes a region-specific reversal in the vector of PCP asymmetry.

**Figure 6.**
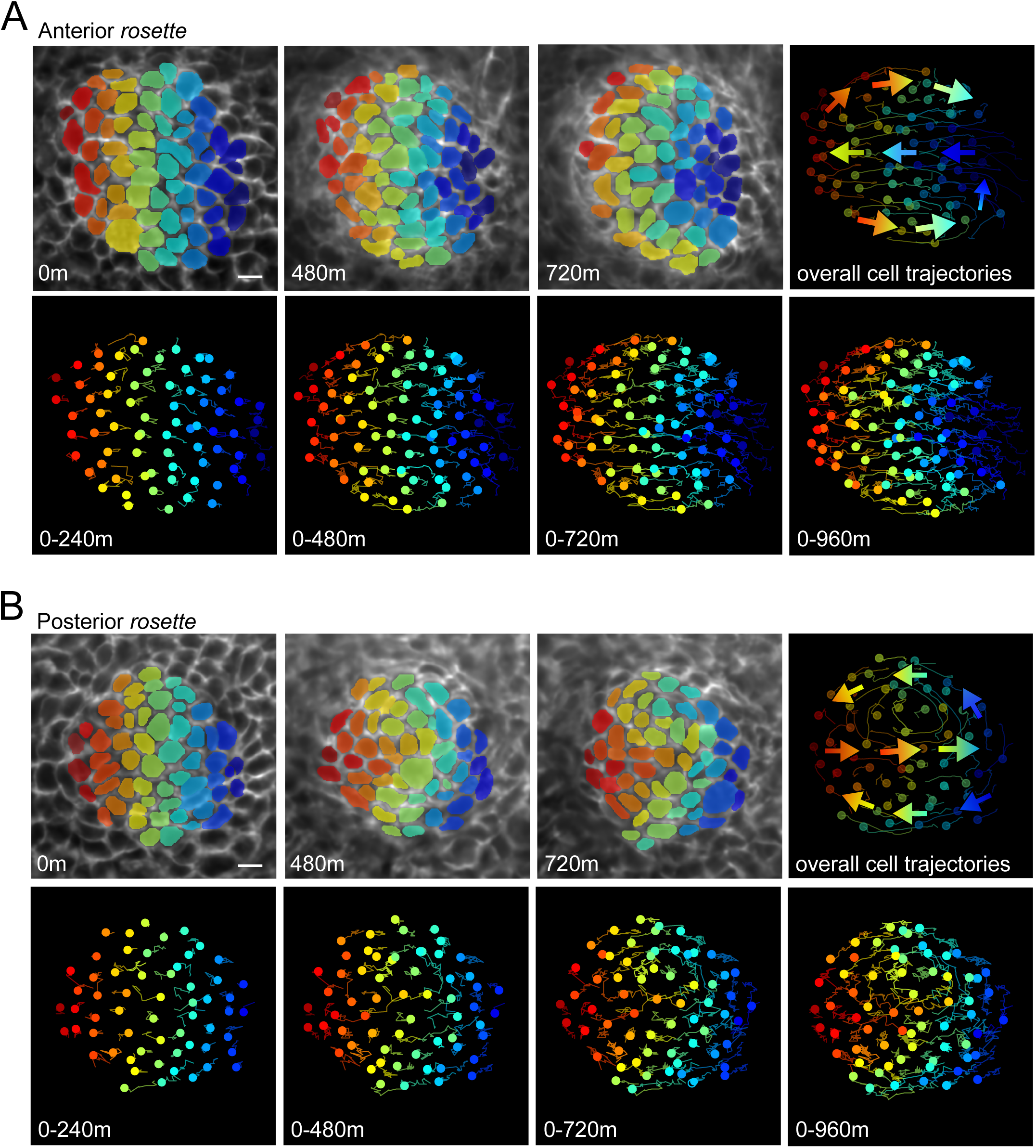
PCP-directed collective cell movements are reversed in posterior *rosette* hair placodes. Spinning disk confocal images from a time series of *rosette* placode cells expressing mTomato (top). The z plane changes to follow the base of the placode into the dermis. Cells were segmented and false colored in a rainbow pattern of vertical lines at t=0. Cell tracks show the movement of cells during the designated time window with circles indicating the last position (bottom). Smoothed tracks with arrow overlays show overall movements through the course of the time series (upper right). (A) Anterior placode, see Supplemental Video 3 and additional example from another explant in Supplemental Video 4. n=7. (B) Posterior placode from the same explant as (A), see Supplemental Video 5 and additional example from another explant in Supplemental Video 6. n=9. Scale bar, 10 µm. Anterior is to the left.

## Discussion

Planar polarized structures typically orient uniformly across entire tissues. In the mammalian skin, collectively oriented hair follicles can align over distances of millimeters to meters, depending on the animal. Previous studies established that the core PCP pathway is central to the development of this organism-scale pattern, as mutations affecting the pathway in mice disrupt follicle orientation across most, if not all, of the skin surface. The *rosette* fancy mouse, a natural variant with region-specific whorls in its fur, offered a unique hair patterning defect with the potential to reveal novel mutations that alter PCP signaling and its coordination across entire tissues. We found the posterior whorls of the *rosette* fancy mouse develop from a 180-degree rotation of the PCP axis which reverses the collective cell rearrangements that orient the hair follicle, revealing that anterior and posterior regions of the skin can establish an axis of polarity independently. We determined the whorled phenotype is associated with a missense mutation in *Fzd6* that alters the consensus sequence required for N-linked glycosylation, trapping the Fzd6 protein in the ER. The Fzd6 mutation is necessary for hair follicle reversal as overexpression of the wild-type protein restores the normal hair follicle pattern. It is unclear, however, how tissue-level mislocalization of Fzd6 can translate into a region-specific rotation of the PCP axis and further studies are required to indicate whether the Fzd6S354I mutation is sufficient to induce reversal.

### The *rosette* phenotype is distinct from other PCP mutants

The *rosette* mouse displays a unique set of PCP phenotypes that are distinct from all previously described mutants that alter epidermal planar polarity, which are comprehensively summarized in Table S4. PCP mutants generally fall into two main categories: complete or partial loss of PCP function. When PCP function is completely lost, epidermal PCP asymmetry is not established, and the directed cell rearrangements that orient embryonic hair placodes are inhibited, causing follicles to grow straight down (Basta et al. 2023; Devenport and Fuchs 2008; Dong et al. 2018; Cetera et al. 2018). Postnatally, the initially vertical follicles rotate to form complex patterns of whorls across the animal’s body, including the rear paws (Cetera et al. 2017; Basta et al. 2023). When PCP activity is only partially compromised, due to functional redundancy as in Fz6 knockouts, less severe epidermal phenotypes arise. Under this condition, PCP proteins also fail to polarize, but cell rearrangements in the hair placode can still occur in an uncoordinated manner to create a pattern of randomly oriented hair follicles (Cetera et al. 2018). In contrast to a complete loss of PCP, residual PCP activity causes misoriented hair follicles to rotate postnatally to correct their orientation along the AP axis, except for paw whorls which are maintained throughout the animal’s life (Wang, Chang, and Nathans 2010; Wang, Badea, and Nathans 2006; Cetera et al. 2017).

The epidermal planar polarity defects exhibited by the *rosette* mouse cannot be fully explained as arising from a complete or partial loss of PCP function when comparing it with previously characterized PCP mutants. Notably, PCP asymmetry is established in embryonic *rosette* skin, but the axis of asymmetry is no longer coordinated across the tissue leading to locally polarized zones that rotate a full 180 degrees in the posterior of the animal. PCP rotation reverses the direction of collective cell movements that orient the follicle resulting in a highly coordinated reversal zone. During early postnatal stages, the reversal zone is maintained, and the posterior whorls emerge. Interestingly, hairs on the rear paws of *rosette* animals are correctly oriented, suggesting PCP activity is not altered in the limbs as it is in other PCP mutants.

A key difference in epidermal polarity defects between *rosette* and PCP mutants is its region-specific reversal. Interestingly, the trafficking defect caused by the Fzd6 S354I mutation does not result in the Fzd6 null pattern of randomly oriented follicles. Further, posterior-specific loss of Fzd6 activity leads to posterior-specific follicle randomization rather than reversal (H. Chang et al. 2016). One possibility is that a small amount of Fzd6 protein, beyond our level of detection, makes it to the membrane resulting in a partial loss of function mutant that is sufficient to organize PCP along the correct axis anteriorly but insufficient to properly coordinate PCP posteriorly. It is unclear, however, how PCP reversal would develop under this condition.

### Naturally occurring zones of polarity reversal

The opposing follicle orientations of the *rosette* mutant is reminiscent of naturally occurring reversal of planar polarized structures in other systems. Sensory hair cells of the mammalian inner ear and neuromasts of the zebrafish display “planar bipolarity” in which hair cell orientation in half of the structure is reflected 180 degrees, creating a mirror image (Kozak et al. 2020; Deans 2021; Tarchini 2021). In both organs, expression of the homeobox transcription factor Emx2 coincides with reversed sensory cell polarity even though the axis of PCP asymmetry is uniform across all cells (Deans et al. 2007; Jiang, Kindt, and Wu 2017; Holley et al. 2010; Kozak et al. 2020; Mirkovic, Pylawka, and Hudspeth 2012; Jacobo et al. 2019). This indicates Emx2 acts downstream of PCP signaling to control how cells transduce directional information into polarized outputs.

The *Drosophila* eye disc presents another example of “planar bipolarity” where ommatidia are reversed at the “equator” located at the dorsal-ventral (DV) midline (Jenny 2010; Koca, Collu, and Mlodzik 2022). In contrast to the previous examples, the vector of PCP asymmetry is also reversed 180 degrees at the equator (Strutt et al. 2002; Rawls and Wolff 2003). Similar to Emx2 expression in the inner ear, homeotic genes are differentially expressed in one half of the eye disc (McNeill et al. 1997; Axelrod and McNeill 2002). However, they do not act by reversing the axis of polarity in a cell-autonomous manner but instead define the position of the equator. Ommatidia polarity can be inverted by inducing ectopic equators or by manipulating DV patterning genes, which are thought to define the axis of PCP asymmetry as upstream directional cues (Tomlinson, Strapps, and Heemskerk 1997; Wehrli and Tomlinson 1998; Zeidler, Perrimon, and Strutt 1999; Cho and Choi 1998; Domínguez and de Celis 1998; Papayannopoulos et al. 1998; C.-H. Yang, Axelrod, and Simon 2002; Rawls, Guinto, and Wolff 2002; McNeill et al. 1997; Maurel-Zaffran and Treisman 2000; Cavodeassi et al. 1999). Taken together, planar bipolarity can be generated in two main ways: through region-specific patterns of gene expression that reverse polarity downstream of the PCP pathway or by reorienting the vector of PCP itself.

Our results suggest that, similar to the organization of PCP in the *Drosophila* eye, hair follicle reversals in *rosette* mutants occur because the vector of PCP in the posterior region of the skin is rotated 180 degrees. Though our data do not directly demonstrate a reversal of core PCP protein asymmetry, we infer the PCP vector is flipped based on the observed rotation of junctional Celsr1 asymmetry in the transition zone where hair follicles reverse. To unambiguously resolve the vector of PCP asymmetry, super-resolution imaging of both Fzd6 and Vangl2 at epithelial cell junctions is required, which is not possible in *rosette* mutants as Fzd6 fails to traffic to the membrane (Basta et al. 2021; Stahley et al. 2021). That hair follicle orientations misalign precisely in the region where Celsr1 asymmetry is observed to rotate strongly suggests it is the axis of PCP itself, and not just the downstream response to PCP signaling, that is responsible for hair follicle reversals in *rosette* mutants.

In the mouse embryo, the Caudal-type homeobox transcription factors, CDX1 and CDX2, are required for AP patterning and posterior axis elongation (van den Akker et al. 2002; Chawengsaksophak et al. 2004). They are expressed in the posterior half of the embryo and therefore are well-positioned to drive posterior-specific gene expression to influence PCP in a regional manner. In fact, CDX genes regulate the expression of core PCP genes *Dvl1* and *Dvl2*, as well as the PCP regulator Ptk7 (Savory et al. 2011). Whether the reversal zone of *rosette* mutants is patterned by CDX targets will be important to determine. The proximal promoter of human CDX2 has been used to alter gene expression specifically in the posterior region of the mouse including the skin (Hinoi et al. 2007; H. Chang et al. 2016; Simonson, Oldham, and Chang 2022). The CDX2 promoter was recently used to drive posterior expression of Wnt5a, which can act as a directional cue to reorient the axis of PCP across several cell diameters (Chu and Sokol 2016; Simonson, Oldham, and Chang 2022). Under this condition, hair follicle orientation was altered specifically in the posterior region of the skin, but follicles were randomized rather than reversed as we observe in *rosette* mutants (Table S4) (Simonson, Oldham, and Chang 2022). Further, Wnt5a knockout mice do not display PCP defects in the skin, and the directional cues that orient epidermal PCP remain to be discovered (Simonson, Oldham, and Chang 2022). Whether or not posterior-specific overexpression of directional cues could reverse the axis of PCP polarity is unknown.

### Epidermal polarity reversal in natural variants

The *rosette* mutant is not the only natural variant that alters epidermal planar polarity in a distinct location. In the crested pigeon, feathers are reversed specifically around the head, whereas in the ridgeback dog, fur orientation is reversed along the posterior midline. The pigeon crest correlates with a point mutation in the kinase domain of EphB2 while the ridgeback trait is linked to a duplication of FGFs 3,4, and 19 (Shapiro et al. 2013; Salmon Hillbertz et al. 2007). Because the mutations alter hair and feather orientation in specific regions while the rest of the skin appears to be unaffected, we reason planar polarity is only altered in a spatially defined zone in these breeds. What determines the precise location of the reversal zone and whether or not the mutations alter the axis of PCP asymmetry, or only the downstream response to PCP, remains to be investigated. The fact that different mutations alter epidermal polarity in distinct regions of the skin suggests that underlying spatial properties determine whether a zone will be refractory or sensitive to a given genetic change.

Altogether, this study provides evidence that the global axis of PCP asymmetry can be regionally decoupled across the epidermis. The missense mutation in Fzd6 present in the *rosette* mouse uncovered an ability of anterior and posterior domains to establish axes of asymmetry in opposing directions leading to a model where a large tissue, such as the skin utilizes multiple inputs at different positions to establish tissue-level patterns.

## Acknowledgements

We gratefully acknowledge those who provided mouse lines, technical support, and valuable discussions that contributed to this project. We thank Mike Chiodo, Bee Bee Mousies, and the fancy mouse breeding community for donating mice and for helpful discussions about the *rosette* line. Sarah Tan developed QuantifyPolarity and provided assistance. We thank Katie Little and Connie Corcoran for assistance with genetic crosses and genotyping, and members of the Devenport lab for insightful comments and suggestions. Finally, we thank Gary Laevsky for imaging support and expertise. The Confocal Facility at Princeton University is a Nikon Center of Excellence. Research reported in this publication was supported by the Eunice Kennedy Shriver National Institute of Child Health and Human Development under award number R00HD097298 to MC and R01HD105009 to DD, the National Science Foundation Graduate Research Fellowship grant number DGE-2039656 to RS, and the National Institute of Arthritis and Musculoskeletal and Skin Diseases of the National Institutes of Health under award number R01AR066070 to DD.

---

Figure S1. **Schematic of regions imaged throughout this study.** (A,B)The transition zone is indicated by the black dotted line. M=medial, L=lateral, A=anterior, P=posterior. (A) Boxes indicate the relative size of imaged regions from e15.5 embryonic flanks in Figures 1B (blue), 1C, 1D, 3E, 3F (black), 4A (arrow), 5B, 5E (magenta), 5C (green), 5D (dotted magenta) 6, S4 (yellow). Multiple positions were imaged on a single flank at different locations in both anterior and posterior regions for Figures 4A, 6, S4. (B) The box indicates the area imaged for P4 pups in Figures 2 and S1. The entire dorsal skin was analyzed. Mouse images were generated by BioRender.

Figure S2. **Coordinated hair follicle reversal persists into postnatal stages.** Additional images of cleared, flat-mounted dorsal skins at postnatal day 4 from *rosette/+* controls (top), *Fzd6-/-* (middle), and *rosette* (bottom). Hair follicle orientation is indicated by the color overlaid on the hair follicle image (cool colors= anteriorly oriented follicles, warm colors= posteriorly oriented follicles). Local order maps corresponding to the same images. Local order is calculated across a radius of 200 pixels containing approximately 200-300 follicles (high order= white, low order= red). AP= anterior-posterior, ML=medial-lateral with M indicating the midline. Scale bar= 1mm.

Figure S3. **The *rosette* phenotype is not due to a naturally occurring deletion in Astrotactin2.** PCR product amplified from exon 5 of Astrotactin2 (Astn2). The expected 89bp product is present in C57BL/6 and *rosette* animals. DNA ladder is Low Molecular Weight DNA Ladder from NEB.

Figure S4. **The N-X-S consensus sequence is required for Fzd6 membrane localization.** (A) Alignment of Fz6 showing conservation of the N-I-S sequence (black rectangle) between mouse, rat, human, chicken, and frog. *Drosophila* Fz lacks the sequence. Amino acids are color coded by their chemistry. (B) Keratinocytes transfected with HA-Fzd6-tdTomato constructs (WT=left, S354I=center, N352A=right) and labeled with anti-HA antibodies with permeabilization. Maximum projections of wide-field stacks are shown. Scale bar, 10 microns.

Figure S5. **The axis of PCP asymmetry rotates in the *rosette* epidermis.** (A) Magnitude of Celsr1 polarity in individual cells within the transition zone of the *rosette* mutant (blue) and corresponding area in controls (green) and Fzd6-/- embryos (magenta). Control n=46,256 cells, *Fzd6-/-* n=45,854 cells, and *rosette* n=48,031 cells from 3 embryos/genotype. (B) Additional examples of long-range polarity patterns shown in Figure 5D.

Figure S6. **PCP-directed collective cell movements occur normally in rst/+ hair placodes.** Spinning disk confocal images from a time series of control *rst/+* placode cells expressing mTomato (top). The z plane changes to follow the base of the placode into the dermis. Cells were segmented and false colored in a rainbow pattern of vertical lines prior to polarization. Cell tracks show the movement of cells during the designated time window with circles indicating the last position (bottom). Smoothed tracks with arrow overlays show overall movements through the course of the time series (upper right). (A) Anterior placode, see Supplemental Video. n=4. (B) Posterior placode from the same explant as (A), See Supplemental Video 2. n=3. Scale bar, 10 µm. Anterior is to the left.

## Videos

Video S1. **PCP-directed collective cell movements occur normally in control explants.** Live imaging of the anterior control placode tracked cells in Figure S4A. Epidermal nuclei are shown in magenta (H2BGFP) and cell membranes are shown in green (mTomato). 17 hours/ 20 minute interval. Scale bar, 10 µm. Anterior is to the left.

Video S2. **PCP-directed collective cell movements occur normally in control explants.** Live imaging of the posterior control placode tracked cells in Figure S4B. Nuclei are shown in magenta (H2BGFP) and cell membranes are shown in green (mTomato). 17 hours/ 20 minute interval. Scale bar, 10 µm. Anterior is to the left.

Video S3. **PCP-directed collective cell movements occur normally in anterior *rosette* placodes.** Live imaging of the anterior *rosette* placode tracked cells in Figure 6A. Nuclei are shown in magenta (H2BGFP) and cell membranes are shown in green (mTomato). 17 hours/ 20 minute interval. Scale bar, 10 µm. Anterior is to the left.

Video S4. **PCP-directed collective cell movements occur normally in anterior *rosette* placodes.** Live imaging of an additional anterior *rosette* placode expressing mTomato to label cell membranes. 17 hours/ 20 minute interval. Scale bar, 10 µm. Anterior is to the left.

Video S5. **PCP-directed collective cell movements are reversed in posterior *rosette* placodes.** Live imaging of the posterior *rosette* placode tracked cells in Figure 6B. Nuclei are shown in magenta (H2BGFP) and cell membranes are shown in green (mTomato). 17 hours/ 20 minute interval. Scale bar, 10 µm. Anterior is to the left.

Video S6. **PCP-directed collective cell movements are reversed in posterior *rosette* placodes.** Live imaging of an additional posterior *rosette* placode expressing mTomato to label cell membranes. 17 hours/ 20 minute interval. Scale bar, 10 µm. Anterior is to the left.

---

**Table S1.** Mouse samples used for SNP analysis scored by phenotype. (Separate file) Whorled animals are represented by a “one” and non-whorled animals are represented by a “zero” in the “Phenotype” column. Familial groups are indicated along with approximate percent C57BL/6 background.

**Table S2.** Summary of mouse sample familial groups and approximate C57BL/6 ancestry. The number of samples from each familial group is indicated under “N.”

| Sibship | Percent. C57BL/6 ancestry | N |
| --- | --- | --- |
| B (Fancy founder) | 0 | 1 |
| B (Sibling of fancy founder; not crossed) | 0 | 1 |
| A | 71.9 | 56 |
| C | 81.3 | 2 |
| D | 56.3 | 4 |
| E | 64.1 | 6 |
| F | 64.1 | 1 |
| G | 64.1 | 5 |
| H | 67.2 | 5 |
| I | 71.9 | 2 |
| J | 71.9 | 2 |
| K | 56.3 | 1 |
| L | 56.3 | 2 |
| Parental animal to A, C, D, H, K, L | 62.5 | 1 |
| C57BL/6 (Lab strain) | 100 | 2 |

**Table S3.**
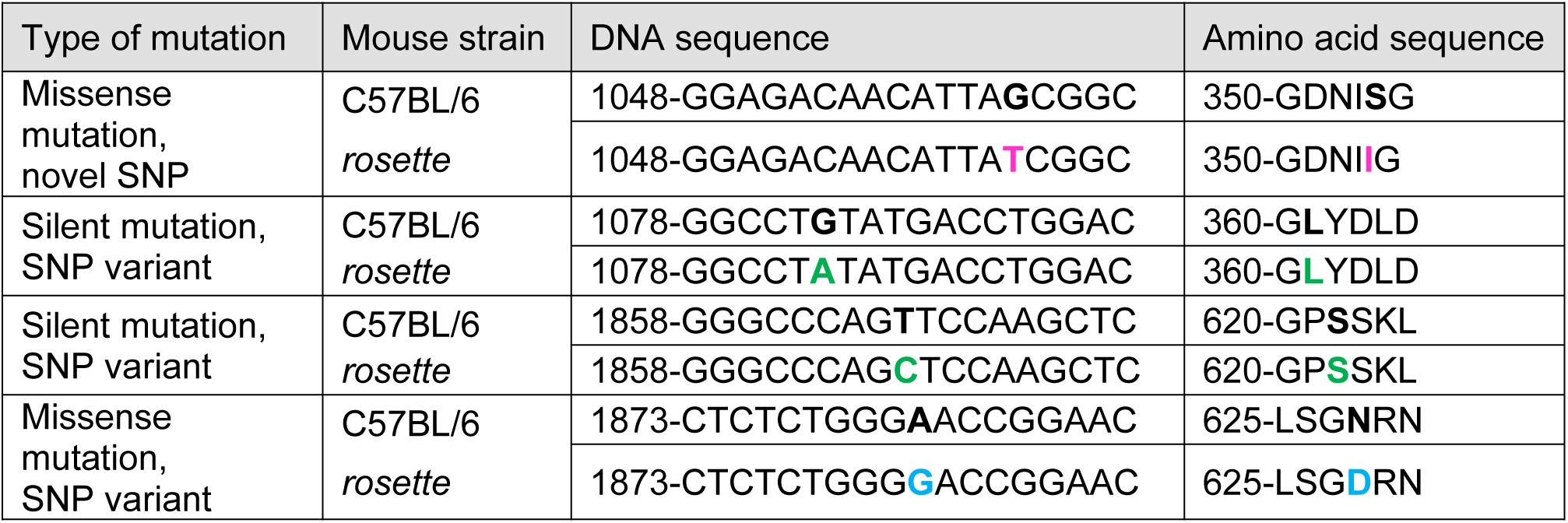
*Fzd6* SNP variants between C57BL/6 and rosette determined by Sanger sequencing. Two mutations are silent (green), one is a known, missense SNP variant (blue), and one is a novel missense mutation (magenta).

**Table S4.**
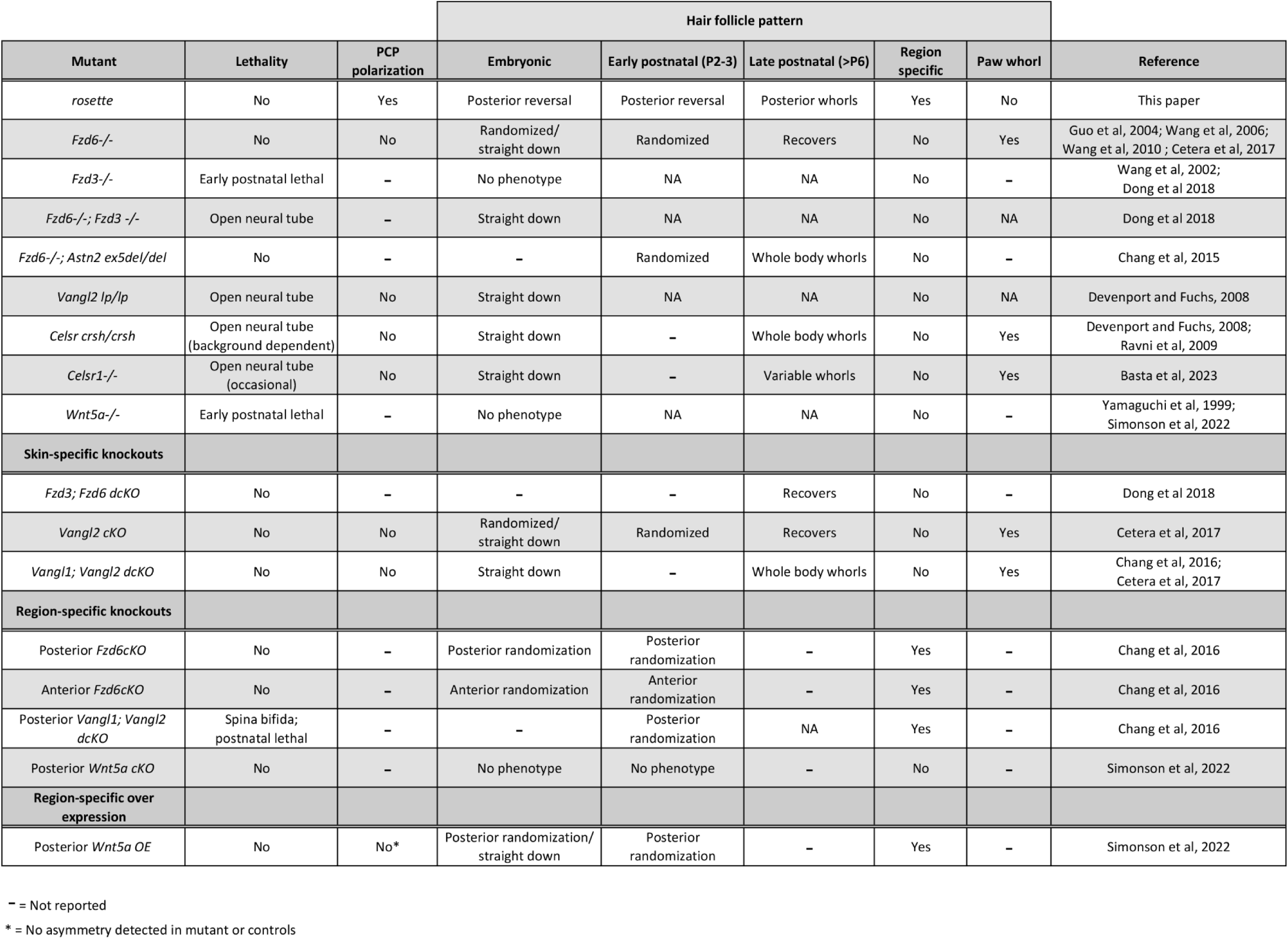
Summary of PCP and *rosette* mutant phenotypes. Comprehensive summary of differences between previously characterized PCP mutants and *rosette*.

**Table S5.** Mouse genotypes.

| Figures | Stages | Genotypes |
| --- | --- | --- |
| **1** | Adult, E15.5 | *C57BL/6*,<br>*Fzd6* *-/-*,<br>*rosette/rosette* |
| **2** | P4 | *rosette/+*,<br>*Fzd6* *-/-* *,<br>*rosette/rosette* |
| **3A-C** | Postnatal | **See Table S1** |
| **3D** | Postnatal | *Unrelated whorled and non-whorled fancy mice* |
| **3E** | E15.5 | *Fzd6* *+/-*,<br>*rosette/+*,<br>*Fzd6* *+/-*; *rosette/+* |
| **3F** | E15.5 | *Rosa26 lox-stop-lox Fzd6WT*; *rosette/rosette*,<br>*K14-Cre*; *Rosa26 lox-stop-lox Fzd6WT*; *rosette/rosette* |
| **4A** | E15.5 | *C57BL/6*,<br>*rosette/rosette* |
| **5** | E15.5 | *C57BL/6*,<br>*Fzd6* *-/-*,<br>*rosette/rosette* |
| **6** | E15.5 | *K14-Cre*; *ROSA26 mTmG*; *K14-H2BGFP/+*; *Fzd6 rosette/+*,<br>*K14-Cre*; *ROSA26 mTmG*; *Fzd6 rosette/+*,<br>*K14-Cre*; *ROSA26 mTmG*; *K14-H2BGFP/+*; *Fzd6 rosette/rosette*,<br>*K14-Cre*; *ROSA26 mTmG*; *Fzd6 rosette/rosette* |

\*P4 *Fzd6* *-/-* whole skin images were reused from Cetera, 2017.

**Table S6.** Mouse allele source.

| Allele | Source |
| --- | --- |
| *C57BL/6J* | JAX 000664 |
| *Fzd6* *-/-* | Guo et al. 2004 (JAX 012824) |
| *rosette/rosette* | Breeding community |
| *Rosa26 lox-stop-lox Fzd6WT* | Hua et al. 2014 (JAX 023843) |
| *K14-Cre* | Vasioukhin et al. 1999 |
| *ROSA26 mTmG* | Muzumdar et al. 2007 (JAX 007676) |
| *K14-H2BGFP* | Tumbar et al. 2004 |

**Table S7.** Antibodies.

| **Antibody** | **Source** | **Concentration** |
| --- | --- | --- |
| Guinea pig anti-Celsr1, polyclonal | D. Devenport (Devenport and Fuchs, 2008) | 1:1000 |
| Rat anti-E-cadherin, monoclonal | ThermoFisher, DECMA-1 clone, 14-3249-82 | 1:1000 |
| Rat anti-P-Cadherin, monoclonal | Clontech, PCD-1 clone, M109 | 1:200 |
| Rabbit anti-Sox9, polyclonal | Millipore, AB5535 | 1:1000 |
| Goat anti-Fzd6, polyclonal | R&D Systems, AF1526 | 1:400 |
| Rat anti-HA, monoclonal | Roche, 3F10 clone, 1867423001 | 1:250-1:500 |
| Hoechst | Invitrogen, H1399 | 1:1000 |
| Donkey anti-guinea pig 647 | Invitrogen, A211450 | 1:1000 |
| Donkey anti-rat 488 | Invitrogen, A21208 | 1:1000 |
| Donkey anti-goat 555 | Invitrogen, A21432 | 1:1000 |
| Donkey anti-rabbit 555 | Invitrogen, A31572 | 1:1000 |
| Donkey anti-rat 647 | Jackson ImmunoResearch, 712605153 | 1:1000-1:2000 |

**Table S8.** Summary of linear mixed model for genetic associations with the binary whorl phenotype. Bolded values indicate a chromosomal enrichment of outlier ß values.

| Chr | N<sub>SNPs</sub> | SNP density (Kb) | N<sub>SNPs</sub> with outlier $\beta$ values | Prop. of chromosome with outlier $\beta$ values |
| --- | --- | --- | --- | --- |
| 1 | 9,698 | 19.8 | 683 | 10 |
| 2 | 1,259 | 38.5 | 628 | 10 |
| 3 | 1,721 | 88.8 | 95 | 1 |
| 4 | 5,281 | 28.4 | 902 | 14 |
| 5 | 6,215 | 23.8 | 786 | 12 |
| 6 | 2,631 | 55.3 | 49 | 1 |
| 7 | 4,788 | 29.7 | 18 | 0 |
| 8 | 5,344 | 23.5 | 64 | 1 |
| 9 | 4,860 | 24.5 | 31 | 0 |
| 10 | 2,571 | 48.8 | 6 | 0 |
| 11 | 1,340 | 86 | 24 | 0 |
| 12 | 2,346 | 46.7 | 108 | 2 |
| 13 | 2,826 | 40.5 | 819 | 13 |
| 14 | 371 | 261.6 | 18 | 0 |
| **15** | **4,976** | **20.2** | **2,092** | **32** |
| 16 | 2,938 | 30.8 | 230 | 4 |
| 17 | 44 | 1,992 | 12 | 0 |
| 18 | 3,792 | 23.1 | 58 | 1 |
| 19 | 662 | 86.6 | 172 | 3 |
| 20 | 1,016 | 157.8 | 0 | 0 |

**Table S9.** Linear mixed model association for 91 mice (85 with known binary whorl phenotypes) with 67K SNPs. (Separate file) Annotations for each SNP is provided, which includes the physical gene name associated, annotated feature, and predicted phenotypic impact. (Abbreviations: AF, allele frequency; L_remle, restricted maximum likelihood estimate of lambda; L_mle, maximum likelihood estimate of lambda; N_miss, number of missing loci; p_LRT, p-value from the likelihood ratio test; S.E., standard errors for ß values).

**Table S10.**
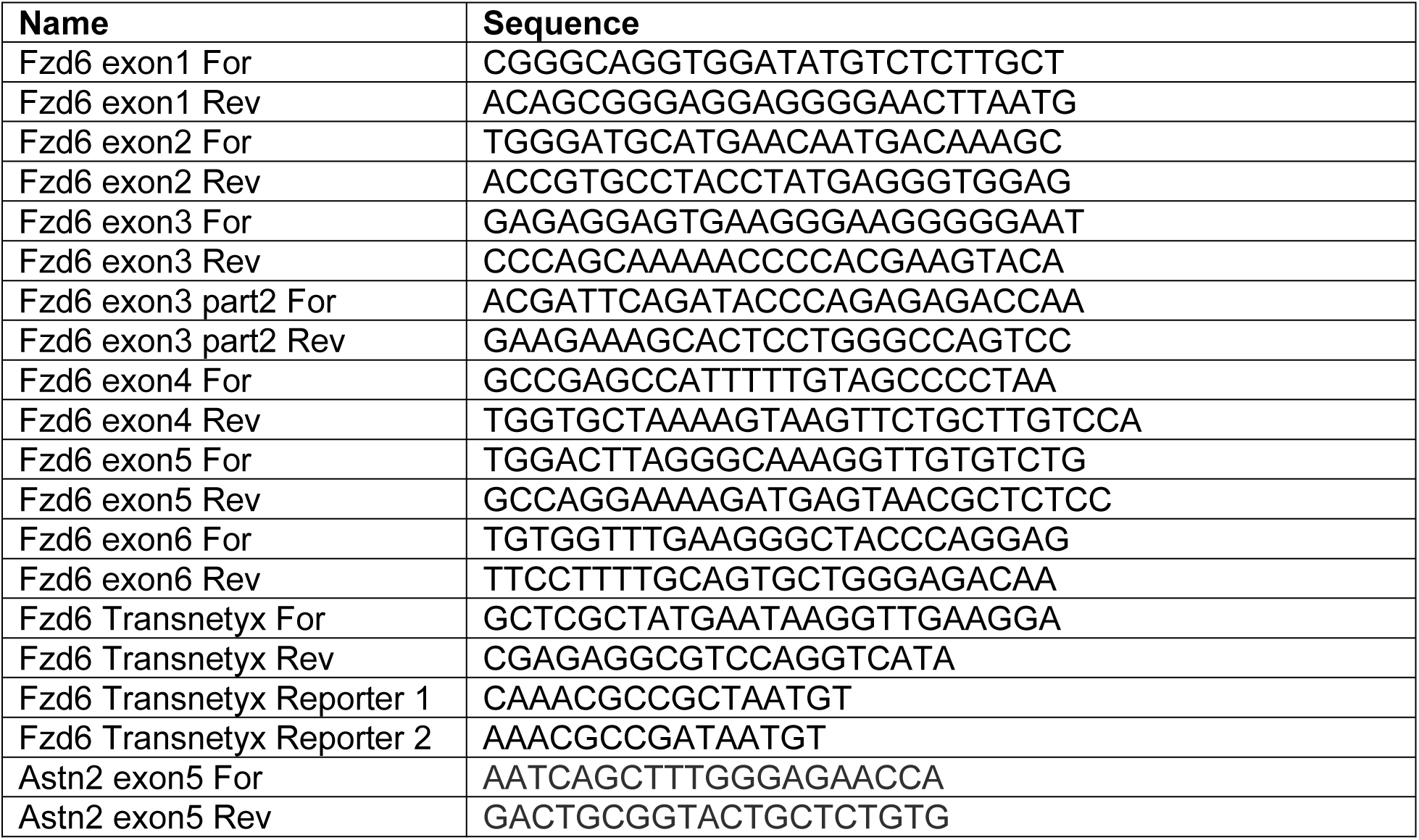
Genotyping primers.

**Table S11.**
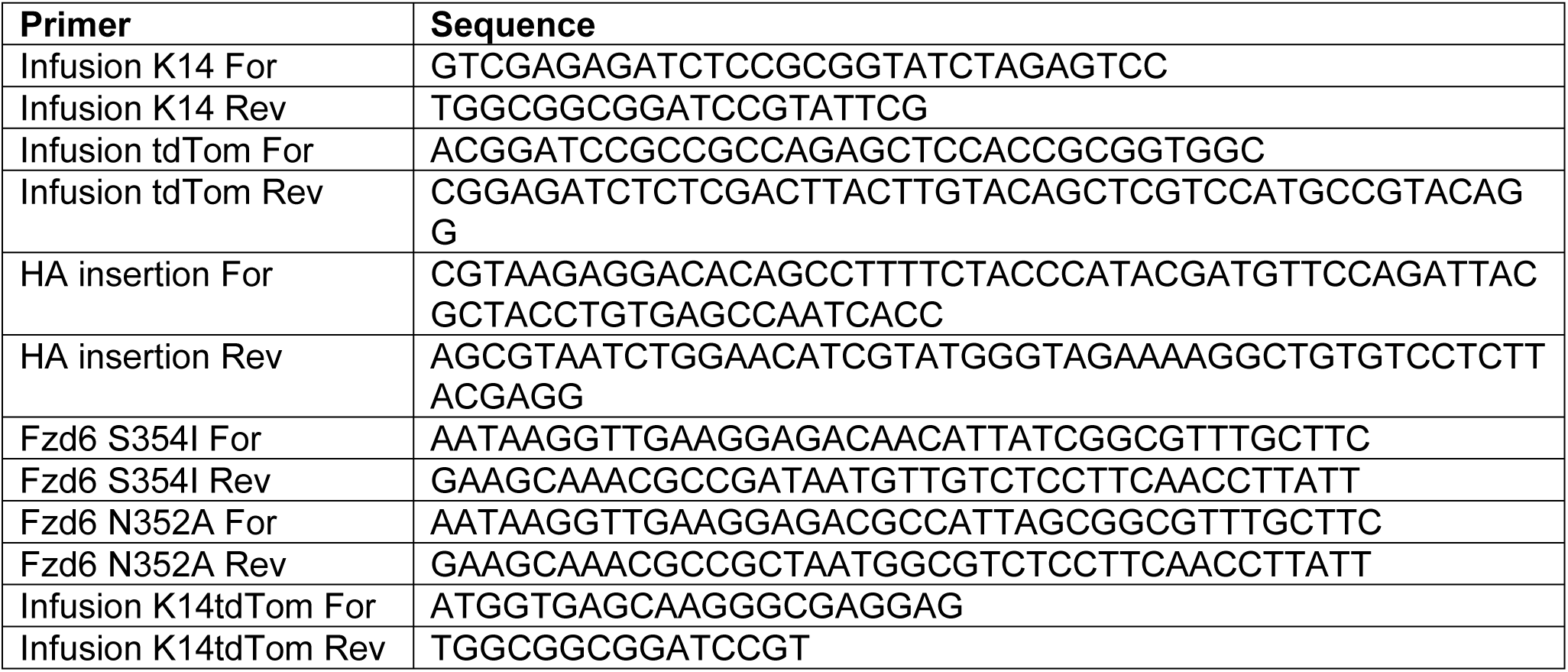
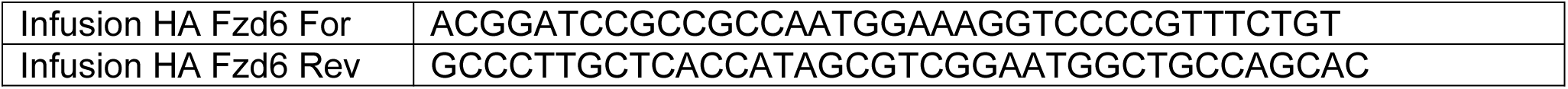
Primers for generating Fzd6 constructs.

## Methods

### Mouse lines and breeding

All procedures involving animals were approved by Princeton University and the University of Minnesota’s Institutional Animal Care and Use Committee (IACUC). Mice were housed in an AAALAC-accredited facility in accordance with the Guide for the Care and Use of Laboratory Animals. Full genotypes are listed in Tables S5, S6. Two male *rosette* animals were donated by a fancy mouse breeder, Mike Chiodo. A single male was used to rederive the line with C57BL/6 females at Charles River. Previously deceased, unrelated whorled and non-whorled fancy animals were donated by BeeBee Mousies.

### Whole-mount immunostaining

E15.5 embryos were dissected in PBS and fixed immediately in 4% paraformaldehyde in DPBS with calcium and magnesium (Gibco cat. 14040) for 1 hour at room temperature. After washing in PBS, back skins were dissected and blocked for 1 hour at room temperature or overnight at 4°C in 3% normal donkey serum, 1% bovine serum albumin, 1% fish gelatin, and 0.02% sodium azide in PBT2 (PBS with 0.2% Triton X-100). Skins were incubated with primary antibodies in blocking solution overnight at 4°C then washed in PBT2. Skins were incubated with secondary antibodies for 2 hours at room temperature or overnight at 4°C, washed, and mounted in Prolong Gold (Molecular Probes). When the P-Cadherin antibody was used, TBS with 0.2% Triton X-100 was used instead of PBT2 for all steps. See Table S7 for antibodies list.

### Quantification of embryonic hair follicle orientation

Entire back skin flanks stained for P-Cadherin and Sox9 were imaged and tiled using a Nikon A1R confocal with a Plan Apo 20x/0.75NA objective. “Focus surface” was used to find a central z-plane at multiple positions across the skin. Images were acquired at the central plane and 5 microns above and below. The ImageJ plugin “3D EDF” was used to flatten the image (Forster et al. 2004). ImageJ and Photoshop were used for image processing.

A semi-automated method was used to detect the location and orientation of hair follicles based on the relative positions of the P-Cadherin and Sox9 labeled cells. This produces an output image with an arrow overlay that can be hand-corrected in ImageJ to fix errors (Basta et al. 2023). The corrected output matrix is used to generate a color-coded Voronoi diagram where each polygon indicates the position of a single follicle and colored according to its orientation.

The order parameter (Equation 1) was calculated by taking the cosine of the difference between two follicle angles (Ɵ0-Ɵi). A local area was determined by taking the average minimum distance between neighboring follicles (d) and multiplying it by 3 (3*d) such that a local area contains a center follicle and the follicles within a 3-follicle radius. This step was repeated for all pairwise combinations in the field and the average was calculated. The output was scaled so the lowest value is 0 and the highest is 1 where 0 represents low order (antiparallel follicles) and 1 represents high order (parallel follicles). The center of the local area was color coded using a heat map (Kovesi 2015) that represents the order parameter.

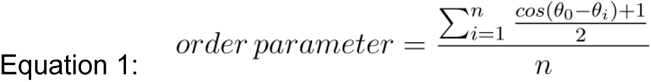

### Sample preparation and quantification of postnatal hair follicle orientation

Skins were processed as previously described (Chang, 2014). After euthanasia, dorsal back skin was dissected from P4 animals and pinned dermis side up to solidified paraffin in a petri dish. Skins were fixed in 4% PFA at 4°C overnight with gentle rocking, then washed with PBS and dehydrated over consecutive days in 70%, 95%, and 100% ethanol. Dehydrated skins were placed in a glass petri dish with a glass slide laid on top to keep the tissue flat. BABB (2:1 ratio of benzyl benzoate and benzyl alcohol) was added to the dish to clear the tissue overnight with gentle rocking at room temperature. Cleared skin was placed between two glass plates for imaging. Brightfield images were acquired on a Nikon SMZ1270 dissecting scope using a Plan Apo 0.5x objective and a Nikon Digital Sight Fi1 camera. Samples were illuminated from beneath the sample using 3-6x magnification. Whole back skin images were obtained by stitching together high magnification images using Photomerge in Photoshop.

The entire dorsal skin was imaged on each animal. To account for differences in animal size from different litters, stitched images were scaled down to match the smallest skin. Thus, the same relative region of each animal was compared. Images were processed using ImageJ. To determine the orientation of each hair follicle, the plugin “OrientationJ-Vector Field” was used (Puspoki, 2016). The local window and grid size was adjusted (10 and 12 for a 2412×1797 pixel image) to reduce noise and allow approximately one vector to represent each follicle. This generated a vector field with the coordinates and orientation of each vector within 180 degrees. Because our phenotypes cause complete hair follicle reversals, we needed to account for 360 degrees. We generated a custom MATLABscript to overlay the arrows on the original image.

We then generated another black and white image overlay where white indicated the vector needed to be flipped (i.e., 0 becomes 180). The updated vector field was then used to generate a color map representing the hair follicle orientation at each position which was overlaid on the original image. The order parameter was calculated as described for the embryonic samples except the local area analyzed was 200 pixels, including approximately 200-300 follicles.

### Genome-wide SNP genotyping array

DNA was extracted from ear punches or tail clips using the Qiagen DNeasy Blood and Tissue kit. We sent high molecular weight DNA to FisherScientific for genotyping array service using the Affymetrix Mouse Diversity Array. The array contains ∼623,000 probe sets that assay single nucleotide polymorphisms (SNPs) previously identified between mouse strains (H. Yang et al. 2009). We used the SNP set to identify SNPs associated with the whorled phenotype. We treated the whorled phenotype as a binary trait and categorized 91 mice (Table S1) as whorled (1) or non-whorled (0). Sample relatedness and ancestry are summarized in Table S2. After the fancy founder was outcrossed to C57BL/6 and backcrossed to phenotypic intermediates, the approximate C57BL/6 background was estimated. The original founder (E09, Table S1) and its sibling that was not used to propagate the line (H05, Table S1) are considered to have a 100% fancy background or 0% C57BL/6. Pure Black6 animals are 100% C57BL/6. We received a filtered data set (541,069 SNP loci) after the vendor performed a quality control step during genotype cluster classification using the Axiom Analysis Suite. “PolyHighResolution,” “NoMinorHom,” “MonoHIghResolution” categories were recommended for further analysis. We applied a minor allele frequency filter (MAF) of <3% using PLINK v1.9 (C. C. Chang et al. 2015) on unique samples that resulted in a final data set of 67,589 SNP loci (67K SNP set). To assess the structuring of the samples with the 67K SNP set, we conducted a clustering analysis using a principle component analysis (PCA) with the program *flashPCA* (Abraham and Inouye 2014) and PC1 captured the gradient of Black6 ancestry.

### Genotype-phenotype association

We used a linear mixed model (*lmm*) in the program *gemma* to identify SNP variants in the 67K set associated with the binary whorl phenotype (present or absent) in the 91 samples (X. Zhou and Stephens 2012). *Gemma* removes SNPs that lack information and ignores individuals that lack phenotypic data. In our sample set, 6/91 samples had an unknown phenotype and were excluded. We constructed a standardized relatedness matrix in *gemma* with the *-gk* 2 function, which was then used in the *lmm* for adjusting association scores for relatedness and used the likelihood ratio test (LRT) for significance testing. We also prioritized SNPs with allelic effect (ß) values in the lower or upper 5^th^-percentile of the distribution and considered these SNPs as putative outliers with outlier ß values. Chromosome 15 contained the highest number of SNPs that were associated with outlier ß values (Table S8). The LRT identified 1,428 SNPs associated with the binary whorl phenotype (*p*<10^-5^), of which all but two were located on mouse chromosome 15 (Table S9). We further identified a candidate region on chromosome 15 (25.7-62.1 Mb) containing 132 genes by integrating the LRT *p*-value and ß values (Figure 3B).

### *Fzd6* genotyping

To sequence all *Fzd6* exons, the primer sets in Table S10 were used for PCR amplification. The product was purified using the Zymo Clean and Concentrate kit and sequenced (Genewiz). To detect the *rosette* mutation, the primer sets for “Fzd6 exon 3 part 2” were used. Ear punches or embryonic tails were also sent to Transnetyx for genotyping where they used the primers and reporters listed in Table S10.

### Frizzled alignment

Using NCBI protein BLAST, the mouse Fzd6 amino acid sequence was aligned to the Fzd6 sequences from rat, human, chicken, frog, and Fz from Drosophila (Altschul et al. 1990). The multiple sequence alignment (MSA) file was downloaded and loaded into RStudio. The ggmsa package was used to display the MSA for extracellular loop 2 with each amino acid colored by its chemistry (Yu 2022; L. Zhou et al. 2022).

### Colocalization of E-Cadherin and Fzd6

E15.5 skins stained for E-Cadherin, Fzd6, and with Hoechst were imaged on an A1R scanning confocal using a Plan Apo 60x/1.4 NA objective. Single z planes in the interfollicular regions were chosen for analysis using only the E-Cadherin and Hoechst channels. In ImageJ, a mask was created using the Auto Threshold/MinError option on the Hoechst channel. This allowed us to exclude the nuclear area in our analysis. Colocalization was measured between the E-Cadherin and Fzd6 channels, excluding the masked area, using the Coloc2 plug-in in ImageJ to determine the Pearson correlation coefficient.

### Molecular cloning and constructs

K14-HA-Fzd6-tdTomato constructs were generated by using In-Fusion (Takara) cloning to first add the tdTomato tag from pBluescript (Stratagene) to the empty K14–β-globin–MCS vector (Vaezi et al. 2002). The HA tag was inserted after the signal sequence of Fzd6 in the pCR2.1-TOPO (Invitrogen) vector by In-Fusion cloning. The S354I and N352A point mutations were introduced to HA-Fzd6 by PCR-mediated site-directed mutagenesis. The final constructs were assembled using In-Fusion cloning to insert the HA-Fzd6 fragments into the K14-tdTomato vector such that the stop sequence of Fzd6 is replaced with tdTomato. The primers used are listed in Table S11.

### Surface detection of HA

CD1 mouse primary keratinocytes were cultured in E-media supplemented with 15% fetal bovine serum and 0.05mM Ca^2+^ at 37°C with 5% CO2 (Nowak and Fuchs 2009). Cells were cultured on fibronectin-treated coverslips and transfected with the K14-HA-Fzd-tdTomato constructs using the Effectene transfection reagent (Qiagen). Cells were switched to high Calcium media (1.5mM Ca^2+^) 12 hours post transfection. Cells were fixed in 4% paraformaldehyde for 10 mins 24 hours post transfection, washed in PBS, incubated for 1 hour with the HA antibody in PBS without detergent. Cells were washed and incubated with a 647 secondary antibody for 30 minutes, washed, and mounted in Prolong Gold.

Stained cells were imaged on a widefield system. Positions were chosen using only the tdTomato signal with the 20x objective. A 60x/1.4NA objective was used to acquire z stacks at the pre-selected positions. For each cell, the exposure time was determined for the tdTomato signal and a scaling factor was determined for HA detection using the WT construct. For example, if a scaling factor of 2 was used for an experiment, then a 200ms exposure time would be used for tdTomato and 400ms for HA on cell 1 and 400ms for tdTomato and 800ms for cell 2. The scaling factor was kept constant for all constructs within an experiment.

Post-processing was performed in ImageJ. Max intensity projections were created for each channel and Auto-thresholding was used to make a mask on the tdTomato channel. The average intensity was calculated within the mask for each channel and the background was subtracted. The ratio of HA signal to tdTomato was calculated for each cell and the average ratio for the WT construct was scaled to 1 to normalize the data. Two cells transfected with the WT construct were omitted from the analysis because the HA signal was oversaturated.

### Spatial representation of PCP protein asymmetry

E15.5 skins stained for Celsr1 and E-Cadherin were imaged on an A1R scanning confocal using a Plan Apo 60x/1.4 NA objective. Z-stacks were imaged and stitched together using Nikon Elements software to generate single images covering over 1mm^2^. To merge z stacks into single in-focus composite images for analysis, 3D EDF (Extended Depth of Field) (Forster et al. 2004) was used in ImageJ on individual channels. Cell masks were created using CellPose (Stringer et al. 2021) or TissueAnalyzer (Aigouy, Umetsu, and Eaton 2016). Celsr1 polarity was measured using QuantifyPolarity which outputs the cell position, magnitude and orientation of Celsr1 asymmetry (Tan et al. 2021). A custom MATLAB script was used to calculate the average vector orientation and magnitude using equations 22 and 23 in (Tan et al. 2021) in a local area with a radius of 40 pixels (center cells and approximately 2 radii of neighbors). The center cell of each neighborhood was color-coded by the average orientation with the saturation indicating the magnitude using a color map (Kovesi 2015). The saturation histogram was scaled such that the bottom and top 1% polarity magnitude values were saturated to 0 and 1 respectively, using the “imadjust” and “stretchlim” functions in MATLAB. Cells with polarity magnitudes less than 0.02 were excluded. The orientation of hair follicles in the area were manually measured using the E-Cadherin channel with no knowledge of Celsr1 asymmetry.

### Live imaging and cell tracking

Live imaging and cell tracking was performed as previously described (Cetera, 2018). E15.5 dorsal skin explants expressing membrane-tdTomato and K14-H2B-GFP were dissected in PBS and transferred onto a 1% agarose gel with F-media containing 10% fetal bovine serum, dermis side down. The agarose gel with the explants was placed into a 35-mm lummox membrane dish (Sarstedt) such that the epidermal surfaces of the explants were in contact with the membrane. Images were acquired on a Nikon Ti-E Spinning Disc with Perfect Focus using a Plan Apo 20/0.75NA air objective and 1.5x optical zoom. Explants were cultured in a humid imaging chamber at 37°C with 5% CO2 during imaging.

The ImageJ plugin MultiStackReg (https://github.com/miura/MultiStackRegistration) was used to correct for XY drift, and a single z plane with the base of the follicle in focus was selected at each time point. Cells were manually tracked in ImageJ using the tdTomato signal and H2B-GFP when present. Cell tracks were produced using the TrackMate plugin (Tinevez et al. 2017). Tracks were then plotted in MATLAB, where tracks were false colored based on their initial AP positions. Overall cell trajectories are smoothed using the ‘smooth’ function in MATLAB with a moving average of 30 time points.

## References

Abraham, Gad, and Michael Inouye. 2014. “Fast Principal Component Analysis of Large-Scale Genome-Wide Data.” PloS One 9 (4): e93766.

Aigouy, Benoit, Daiki Umetsu, and Suzanne Eaton. 2016. “Segmentation and Quantitative Analysis of Epithelial Tissues.” Methods in Molecular Biology 1478: 227–39.

Akker, Eric van den, Sylvie Forlani, Kallayanee Chawengsaksophak, Wim de Graaff, Felix Beck, Barbara I. Meyer, and Jacqueline Deschamps. 2002. “Cdx1 and Cdx2 Have Overlapping Functions in Anteroposterior Patterning and Posterior Axis Elongation.” Development 129 (9): 2181–93.

Altschul, S. F., W. Gish, W. Miller, E. W. Myers, and D. J. Lipman. 1990. “Basic Local Alignment Search Tool.” Journal of Molecular Biology 215 (3): 403–10.

Axelrod, Jeffrey D., and Helen McNeill. 2002. “Coupling Planar Cell Polarity Signaling to Morphogenesis.” TheScientificWorldJournal 2 (February): 434–54.

Basta, Lena P., Michael Hill-Oliva, Sarah V. Paramore, Rishabh Sharan, Audrey Goh, Abhishek Biswas, Marvin Cortez, Katherine A. Little, Eszter Posfai, and Danelle Devenport. 2021. “New Mouse Models for High Resolution and Live Imaging of Planar Cell Polarity Proteins *in Vivo*.” Development. https://doi.org/10.1242/dev.199695.

Basta, Lena P., Parijat Sil, Rebecca A. Jones, Katherine A. Little, Gabriela Hayward-Lara, and Danelle Devenport. 2023. “Celsr1 and Celsr2 Exhibit Distinct Adhesive Interactions and Contributions to Planar Cell Polarity.” Frontiers in Cell and Developmental Biology. https://doi.org/10.3389/fcell.2022.1064907.

Butler, Mitchell T., and John B. Wallingford. 2017. “Planar Cell Polarity in Development and Disease.” Nature Reviews. Molecular Cell Biology 18 (6): 375–88.

Castle, William Ernest. 1905. Heredity of Coat Characters in Guinea-Pigs and Rabbits. Vol. 23. Carnegie Institution of Washington.

Cavodeassi, F., R. Diez Del Corral, S. Campuzano, and M. Domínguez. 1999. “Compartments and Organising Boundaries in the Drosophila Eye: The Role of the Homeodomain Iroquois Proteins.” Development 126 (22): 4933–42.

Cetera, Maureen, Liliya Leybova, Bradley Joyce, and Danelle Devenport. 2018. “Counter-Rotational Cell Flows Drive Morphological and Cell Fate Asymmetries in Mammalian Hair Follicles.” Nature Cell Biology 20 (5): 541–52.

Cetera, Maureen, Liliya Leybova, Frank W. Woo, Michael Deans, and Danelle Devenport. 2017. “Planar Cell Polarity-Dependent and Independent Functions in the Emergence of Tissue-Scale Hair Follicle Patterns.” Developmental Biology 428 (1): 188–203.

Chang, Christopher C., Carson C. Chow, Laurent Cam Tellier, Shashaank Vattikuti, Shaun M. Purcell, and James J. Lee. 2015. “Second-Generation PLINK: Rising to the Challenge of Larger and Richer Datasets.” GigaScience 4 (February): 7.

Chang, Hao, Hugh Cahill, Philip M. Smallwood, Yanshu Wang, and Jeremy Nathans. 2015. “Identification of Astrotactin2 as a Genetic Modifier That Regulates the Global Orientation of Mammalian Hair Follicles.” PLoS Genetics 11 (9): e1005532.

Chang, Hao, Philip M. Smallwood, John Williams, and Jeremy Nathans. 2016. “The Spatio-Temporal Domains of Frizzled6 Action in Planar Polarity Control of Hair Follicle Orientation.” Developmental Biology 409 (1): 181–93.

Chawengsaksophak, Kallayanee, Wim de Graaff, Janet Rossant, Jacqueline Deschamps, and Felix Beck. 2004. “Cdx2 Is Essential for Axial Elongation in Mouse Development.” Proceedings of the National Academy of Sciences of the United States of America 101 (20): 7641–45.

Cho, K. O., and K. W. Choi. 1998. “Fringe Is Essential for Mirror Symmetry and Morphogenesis in the Drosophila Eye.” Nature 396 (6708): 272–76.

Chu, Chih-Wen, and Sergei Y. Sokol. 2016. “Wnt Proteins Can Direct Planar Cell Polarity in Vertebrate Ectoderm.” ELife 5 (September). https://doi.org/10.7554/eLife.16463.

Curtin, John A., Elizabeth Quint, Vicky Tsipouri, Ruth M. Arkell, Bruce Cattanach, Andrew J. Copp, Deborah J. Henderson, et al. 2003. “Mutation of Celsr1 Disrupts Planar Polarity of Inner Ear Hair Cells and Causes Severe Neural Tube Defects in the Mouse.” Current Biology: CB 13 (13): 1129–33.

Davey, Crystal F., and Cecilia B. Moens. 2017. “Planar Cell Polarity in Moving Cells: Think Globally, Act Locally.” Development 144 (2): 187–200.

Deans, Michael R. 2021. “Conserved and Divergent Principles of Planar Polarity Revealed by Hair Cell Development and Function.” Frontiers in Neuroscience 15 (October): 742391.

Deans, Michael R., Dragana Antic, Kaye Suyama, Matthew P. Scott, Jeffrey D. Axelrod, and Lisa V. Goodrich. 2007. “Asymmetric Distribution of Prickle-like 2 Reveals an Early Underlying Polarization of Vestibular Sensory Epithelia in the Inner Ear.” The Journal of Neuroscience: The Official Journal of the Society for Neuroscience 27 (12): 3139–47.

Devenport, Danelle, and Elaine Fuchs. 2008. “Planar Polarization in Embryonic Epidermis Orchestrates Global Asymmetric Morphogenesis of Hair Follicles.” Nature Cell Biology 10 (11): 1257–68.

Domínguez, M., and J. F. de Celis. 1998. “A Dorsal/Ventral Boundary Established by Notch Controls Growth and Polarity in the Drosophila Eye.” Nature 396 (6708): 276–78.

Dong, Bo, Samantha Vold, Cristina Olvera-Jaramillo, and Hao Chang. 2018. “Functional Redundancy of Frizzled 3 and Frizzled 6 in Planar Cell Polarity Control of Mouse Hair Follicles.” Development 145 (19). https://doi.org/10.1242/dev.168468.

Fisher, Katherine H., and David Strutt. 2019. “A Theoretical Framework for Planar Polarity Establishment through Interpretation of Graded Cues by Molecular Bridges.” Development 146 (3). https://doi.org/10.1242/dev.168955.

Forster, B., D. Van De Ville, J. Berent, D. Sage, and M. Unser. 2004. “Extended Depth-of-Focus for Multi-Channel Microscopy Images: A Complex Wavelet Approach.” In 2004 2nd IEEE International Symposium on Biomedical Imaging: Nano to Macro (IEEE Cat No. 04EX821), 660–663 Vol. 1.

Gubb, D., and A. García-Bellido. 1982. “A Genetic Analysis of the Determination of Cuticular Polarity during Development in Drosophila Melanogaster.” Journal of Embryology and Experimental Morphology 68 (April): 37–57.

Guo, Nini, Charles Hawkins, and Jeremy Nathans. 2004. “Frizzled6 Controls Hair Patterning in Mice.” Proceedings of the National Academy of Sciences of the United States of America 101 (25): 9277–81.

Hinoi, Takao, Aytekin Akyol, Brian K. Theisen, David O. Ferguson, Joel K. Greenson, Bart O. Williams, Kathleen R. Cho, and Eric R. Fearon. 2007. “Mouse Model of Colonic Adenoma-Carcinoma Progression Based on Somatic Apc Inactivation.” Cancer Research 67 (20): 9721–30.

Holley, Matthew, Charlotte Rhodes, Adam Kneebone, Michel K. Herde, Michelle Fleming, and Karen P. Steel. 2010. “Emx2 and Early Hair Cell Development in the Mouse Inner Ear.” Developmental Biology 340 (2): 547–56.

Hua, Zhong L., Hao Chang, Yanshu Wang, Philip M. Smallwood, and Jeremy Nathans. 2014. “Partial Interchangeability of Fz3 and Fz6 in Tissue Polarity Signaling for Epithelial Orientation and Axon Growth and Guidance.” Development 141 (20): 3944–54.

Jacobo, Adrian, Agnik Dasgupta, Anna Erzberger, Kimberly Siletti, and A. J. Hudspeth. 2019. “Notch-Mediated Determination of Hair-Bundle Polarity in Mechanosensory Hair Cells of the Zebrafish Lateral Line.” Current Biology: CB 29 (21): 3579–3587.e7.

Jenny, Andreas. 2010. “Planar Cell Polarity Signaling in the Drosophila Eye.” Current Topics in Developmental Biology 93: 189–227.

Jiang, Tao, Katie Kindt, and Doris K. Wu. 2017. “Transcription Factor Emx2 Controls Stereociliary Bundle Orientation of Sensory Hair Cells.” ELife 6 (March). https://doi.org/10.7554/eLife.23661.

Kibar, Zoha, Kyle J. Vogan, Normand Groulx, Monica J. Justice, D. Alan Underhill, and Philippe Gros. 2001. “Ltap, a Mammalian Homolog of Drosophila Strabismus/Van Gogh, Is Altered in the Mouse Neural Tube Mutant Loop-Tail.” Nature Genetics. https://doi.org/10.1038/90081.

King, Robert. 1975. Handbook of Genetics: Volume 4 Vertebrates of Genetic Interest. Vol. 4. Plenum Press, New York.

Koca, Yildiz, Giovanna M. Collu, and Marek Mlodzik. 2022. “Wnt-Frizzled Planar Cell Polarity Signaling in the Regulation of Cell Motility.” Current Topics in Developmental Biology 150 (May): 255–97.

Kovesi, Peter. 2015. “Good Colour Maps: How to Design Them.” ArXiv [Cs.GR]. arXiv. http://arxiv.org/abs/1509.03700.

Kozak, Eva L., Subarna Palit, Jerónimo R. Miranda-Rodríguez, Aleksandar Janjic, Anika Böttcher, Heiko Lickert, Wolfgang Enard, Fabian J. Theis, and Hernán López-Schier. 2020. “Epithelial Planar Bipolarity Emerges from Notch-Mediated Asymmetric Inhibition of Emx2.” Current Biology: CB 30 (6): 1142–1151.e6.

Krasnow, R. E., L. L. Wong, and P. N. Adler. 1995. “Dishevelled Is a Component of the Frizzled Signaling Pathway in Drosophila.” Development 121 (12): 4095–4102.

Lilue, Jingtao, Anthony G. Doran, Ian T. Fiddes, Monica Abrudan, Joel Armstrong, Ruth Bennett, William Chow, et al. 2018. “Sixteen Diverse Laboratory Mouse Reference Genomes Define Strain-Specific Haplotypes and Novel Functional Loci.” Nature Genetics 50 (11): 1574–83.

Maurel-Zaffran, C., and J. E. Treisman. 2000. “Pannier Acts Upstream of Wingless to Direct Dorsal Eye Disc Development in Drosophila.” Development 127 (5): 1007–16.

McNeill, H., C. H. Yang, M. Brodsky, J. Ungos, and M. A. Simon. 1997. “Mirror Encodes a Novel PBX-Class Homeoprotein That Functions in the Definition of the Dorsal-Ventral Border in the Drosophila Eye.” Genes & Development 11 (8): 1073–82.

Mirkovic, Ivana, Serhiy Pylawka, and A. J. Hudspeth. 2012. “Rearrangements between Differentiating Hair Cells Coordinate Planar Polarity and the Establishment of Mirror Symmetry in Lateral-Line Neuromasts.” Biology Open 1 (5): 498–505.

Murdoch, J. N., K. Doudney, C. Paternotte, A. J. Copp, and P. Stanier. 2001. “Severe Neural Tube Defects in the Loop-Tail Mouse Result from Mutation of Lpp1, a Novel Gene Involved in Floor Plate Specification.” Human Molecular Genetics 10 (22): 2593–2601.

Muzumdar, Mandar Deepak, Bosiljka Tasic, Kazunari Miyamichi, Ling Li, and Liqun Luo. 2007. “A Global Double-Fluorescent Cre Reporter Mouse.” Genesis. https://doi.org/10.1002/dvg.20335.

Nowak, Jonathan A., and Elaine Fuchs. 2009. “Isolation and Culture of Epithelial Stem Cells.” In Stem Cells in Regenerative Medicine, edited by Julie Audet and William L. Stanford, 215–32. Totowa, NJ: Humana Press.

Papayannopoulos, V., A. Tomlinson, V. M. Panin, C. Rauskolb, and K. D. Irvine. 1998. “Dorsal-Ventral Signaling in the Drosophila Eye.” Science 281 (5385): 2031–34.

Phifer-Rixey, Megan, and Michael W. Nachman. 2015. “Insights into Mammalian Biology from the Wild House Mouse Mus Musculus.” ELife 4 (April). https://doi.org/10.7554/eLife.05959.

Ravni, Aurélia, Yibo Qu, André M. Goffinet, and Fadel Tissir. 2009. “Planar Cell Polarity Cadherin Celsr1 Regulates Skin Hair Patterning in the Mouse.” The Journal of Investigative Dermatology 129 (10): 2507–9.

Rawls, Amy S., Jake B. Guinto, and Tanya Wolff. 2002. “The Cadherins Fat and Dachsous Regulate Dorsal/Ventral Signaling in the Drosophila Eye.” Current Biology: CB 12 (12): 1021–26.

Rawls, Amy S., and Tanya Wolff. 2003. “Strabismus Requires Flamingo and Prickle Function to Regulate Tissue Polarity in the Drosophila Eye.” Development 130 (9): 1877–87.

Salmon Hillbertz, Nicolette H. C., Magnus Isaksson, Elinor K. Karlsson, Eva Hellmén, Gerli Rosengren Pielberg, Peter Savolainen, Claire M. Wade, et al. 2007. “Duplication of FGF3, FGF4, FGF19 and ORAOV1 Causes Hair Ridge and Predisposition to Dermoid Sinus in Ridgeback Dogs.” Nature Genetics 39 (11): 1318–20.

Savory, Joanne G. A., Melissa Mansfield, Filippo M. Rijli, and David Lohnes. 2011. “Cdx Mediates Neural Tube Closure through Transcriptional Regulation of the Planar Cell Polarity Gene Ptk7.” Development 138 (7): 1361–70.

Shapiro, Michael D., Zev Kronenberg, Cai Li, Eric T. Domyan, Hailin Pan, Michael Campbell, Hao Tan, et al. 2013. “Genomic Diversity and Evolution of the Head Crest in the Rock Pigeon.” Science 339 (6123): 1063–67.

Simonson, Laura, Ethan Oldham, and Hao Chang. 2022. “Overactive Wnt5a Signaling Disrupts Hair Follicle Polarity during Mouse Skin Development.” Development 149 (22). https://doi.org/10.1242/dev.200816.

Stahley, Sara N., Lena P. Basta, Rishabh Sharan, and Danelle Devenport. 2021. “Celsr1 Adhesive Interactions Mediate the Asymmetric Organization of Planar Polarity Complexes.” ELife 10 (February). https://doi.org/10.7554/eLife.62097.

Stringer, Carsen, Tim Wang, Michalis Michaelos, and Marius Pachitariu. 2021. “Cellpose: A Generalist Algorithm for Cellular Segmentation.” Nature Methods 18 (1): 100–106.

Strutt, David, Ruth Johnson, Katherine Cooper, and Sarah Bray. 2002. “Asymmetric Localization of Frizzled and the Determination of Notch-Dependent Cell Fate in the Drosophila Eye.” Current Biology: CB 12 (10): 813–24.

Tan, Su Ee, Weijie Tan, Katherine Fisher, and David Strutt. 2021. “QuantifyPolarity, a New Tool-Kit for Measuring Planar Polarized Protein Distributions and Cell Properties in Developing Tissues.” *Development*, August. https://doi.org/10.1242/dev.198952.

Tang, Xiao, Lina Zhang, Tianji Ma, Mo Wang, Baiying Li, Liwen Jiang, Yan Yan, and Yusong Guo. 2020. “Molecular Mechanisms That Regulate Export of the Planar Cell-Polarity Protein Frizzled-6 out of the Endoplasmic Reticulum.” The Journal of Biological Chemistry 295 (27): 8972–87.

Tarchini, Basile. 2021. “A Reversal in Hair Cell Orientation Organizes Both the Auditory and Vestibular Organs.” Frontiers in Neuroscience 15 (September): 695914.

Taylor, J., N. Abramova, J. Charlton, and P. N. Adler. 1998. “Van Gogh: A New Drosophila Tissue Polarity Gene.” Genetics 150 (1): 199–210.

Tinevez, Jean-Yves, Nick Perry, Johannes Schindelin, Genevieve M. Hoopes, Gregory D. Reynolds, Emmanuel Laplantine, Sebastian Y. Bednarek, Spencer L. Shorte, and Kevin W. Eliceiri. 2017. “TrackMate: An Open and Extensible Platform for Single-Particle Tracking.” Methods 115 (February): 80–90.

Tomlinson, A., W. R. Strapps, and J. Heemskerk. 1997. “Linking Frizzled and Wnt Signaling in Drosophila Development.” Development 124 (22): 4515–21.

Tumbar, Tudorita, Geraldine Guasch, Valentina Greco, Cedric Blanpain, William E. Lowry, Michael Rendl, and Elaine Fuchs. 2004. “Defining the Epithelial Stem Cell Niche in Skin.” Science. https://doi.org/10.1126/science.1092436.

Vaezi, Alec, Christoph Bauer, Valeri Vasioukhin, and Elaine Fuchs. 2002. “Actin Cable Dynamics and Rho/Rock Orchestrate a Polarized Cytoskeletal Architecture in the Early Steps of Assembling a Stratified Epithelium.” Developmental Cell 3 (3): 367–81.

Vasioukhin, V., L. Degenstein, B. Wise, and E. Fuchs. 1999. “The Magical Touch: Genome Targeting in Epidermal Stem Cells Induced by Tamoxifen Application to Mouse Skin.” Proceedings of the National Academy of Sciences of the United States of America 96 (15): 8551–56.

Wallace, M. E. 1971. Mouse News Letters 44: 18.

Wang, Yanshu, Tudor Badea, and Jeremy Nathans. 2006. “Order from Disorder: Self-Organization in Mammalian Hair Patterning.” Proceedings of the National Academy of Sciences of the United States of America 103 (52): 19800–805.

Wang, Yanshu, Hao Chang, and Jeremy Nathans. 2010. “When Whorls Collide: The Development of Hair Patterns in Frizzled 6 Mutant Mice.” Development 137 (23): 4091–99.

Wang, Yanshu, Nupur Thekdi, Philip M. Smallwood, Jennifer P. Macke, and Jeremy Nathans. 2002. “Frizzled-3 Is Required for the Development of Major Fiber Tracts in the Rostral CNS.” The Journal of Neuroscience: The Official Journal of the Society for Neuroscience 22 (19): 8563–73.

Wehrli, M., and A. Tomlinson. 1998. “Independent Regulation of Anterior/Posterior and Equatorial/Polar Polarity in the Drosophila Eye; Evidence for the Involvement of Wnt Signaling in the Equatorial/Polar Axis.” Development 125 (8): 1421–32.

Wolff, T., and G. M. Rubin. 1998. “Strabismus, a Novel Gene That Regulates Tissue Polarity and Cell Fate Decisions in Drosophila.” Development 125 (6): 1149–59.

Yamaguchi, Terry P., Allan Bradley, Andrew P. McMahon, and Steven Jones. 1999. “A *Wnt5a* Pathway Underlies Outgrowth of Multiple Structures in the Vertebrate Embryo.” Development. https://doi.org/10.1242/dev.126.6.1211.

Yang, Chung-Hui, Jeffrey D. Axelrod, and Michael A. Simon. 2002. “Regulation of Frizzled by Fat-like Cadherins during Planar Polarity Signaling in the Drosophila Compound Eye.” Cell 108 (5): 675–88.

Yang, Hyuna, Yueming Ding, Lucie N. Hutchins, Jin Szatkiewicz, Timothy A. Bell, Beverly J. Paigen, Joel H. Graber, Fernando Pardo-Manuel de Villena, and Gary A. Churchill. 2009. “A Customized and Versatile High-Density Genotyping Array for the Mouse.” Nature Methods 6 (9): 663–66.

Yu, Guangchuang. 2022. Data Integration, Manipulation and Visualization of Phylogenetic Trees. CRC Press.

Zeidler, M. P., N. Perrimon, and D. I. Strutt. 1999. “Polarity Determination in the Drosophila Eye: A Novel Role for Unpaired and JAK/STAT Signaling.” Genes & Development 13 (10): 1342–53.

Zhou, Lang, Tingze Feng, Shuangbin Xu, Fangluan Gao, Tommy T. Lam, Qianwen Wang, Tianzhi Wu, et al. 2022. “Ggmsa: A Visual Exploration Tool for Multiple Sequence Alignment and Associated Data.” Briefings in Bioinformatics 23 (4). https://doi.org/10.1093/bib/bbac222.

Zhou, Xiang, and Matthew Stephens. 2012. “Genome-Wide Efficient Mixed-Model Analysis for Association Studies.” Nature Genetics 44 (7): 821–24.

