## Supplemental Figures for "Region-specific reversal of epidermal planar polarity in the fancy *rosette* mouse"

A

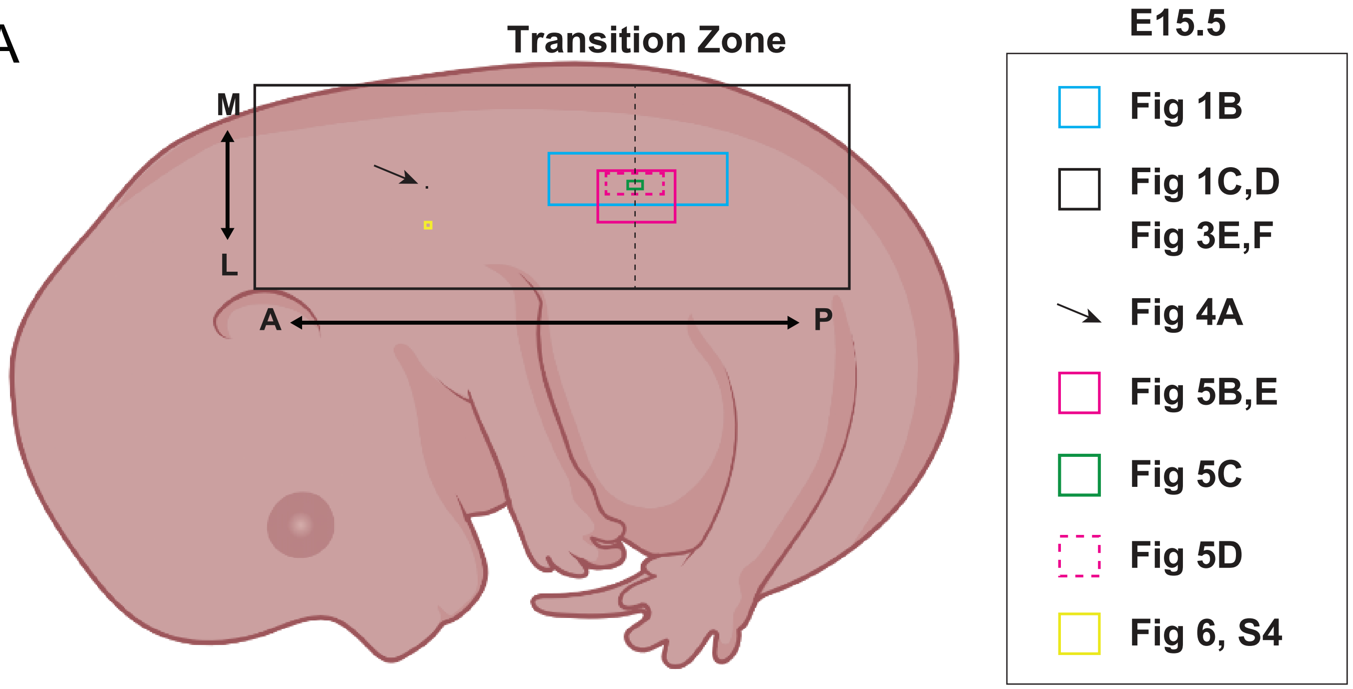


B

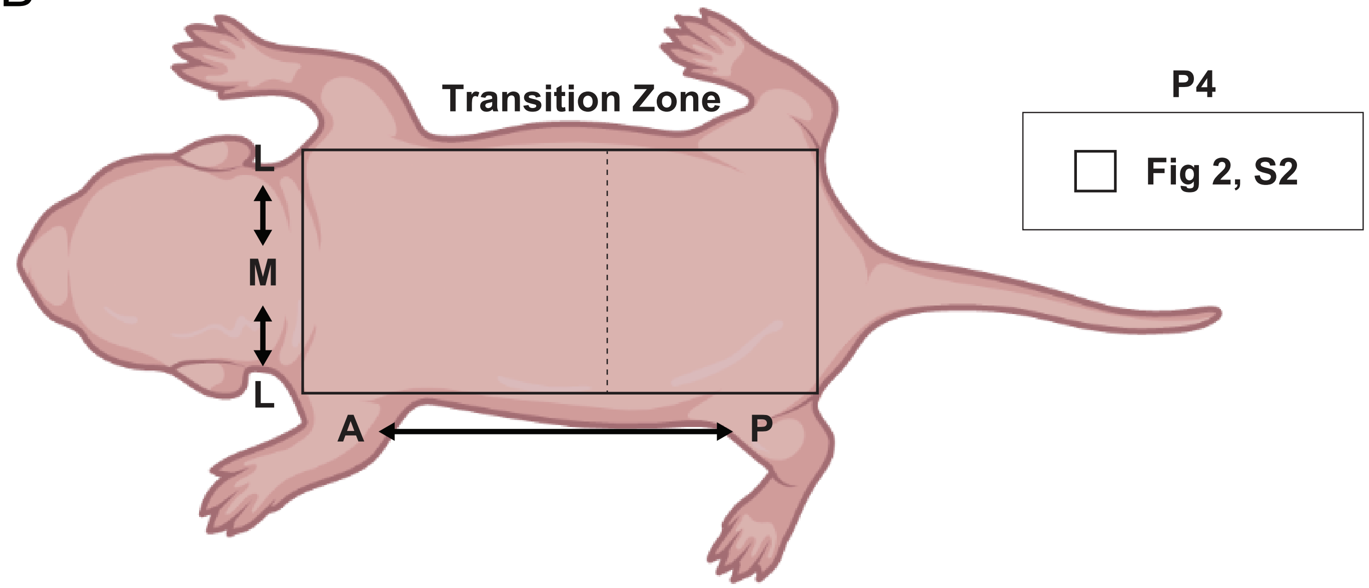


Figure S1. **Schematic of regions imaged throughout this study.**

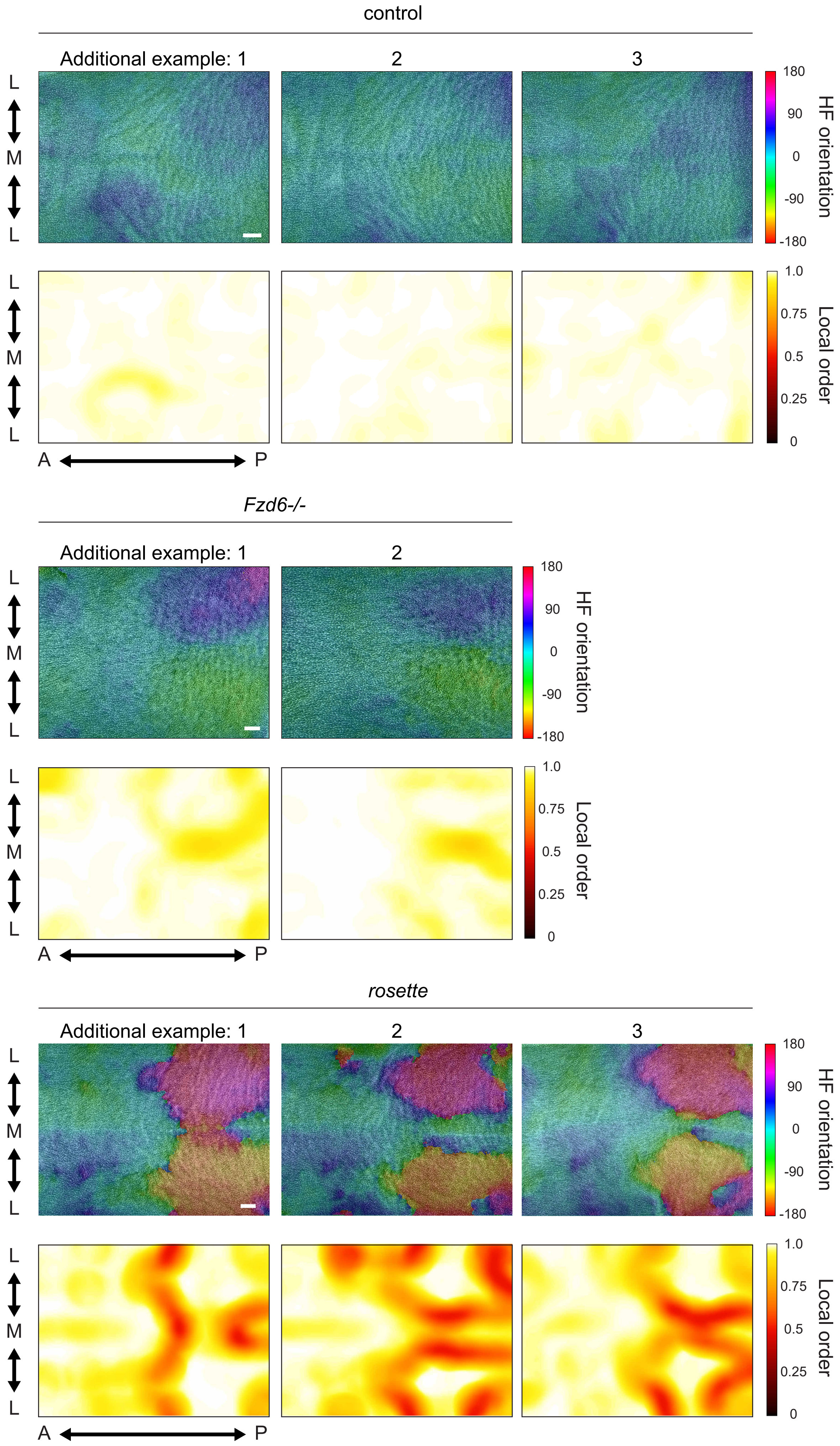


Figure S2. **Coordinated hair follicle reversal persists into postnatal stages.**

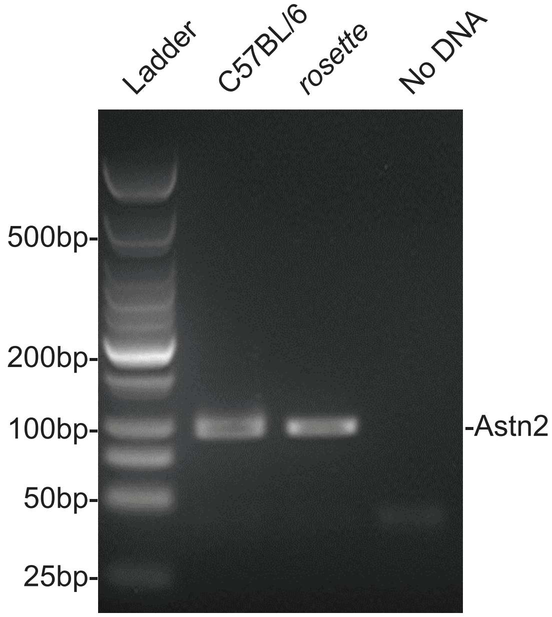


Figure S3. **The *rosette* phenotype is not due to a naturally occurring deletion in Astrotactin2.**

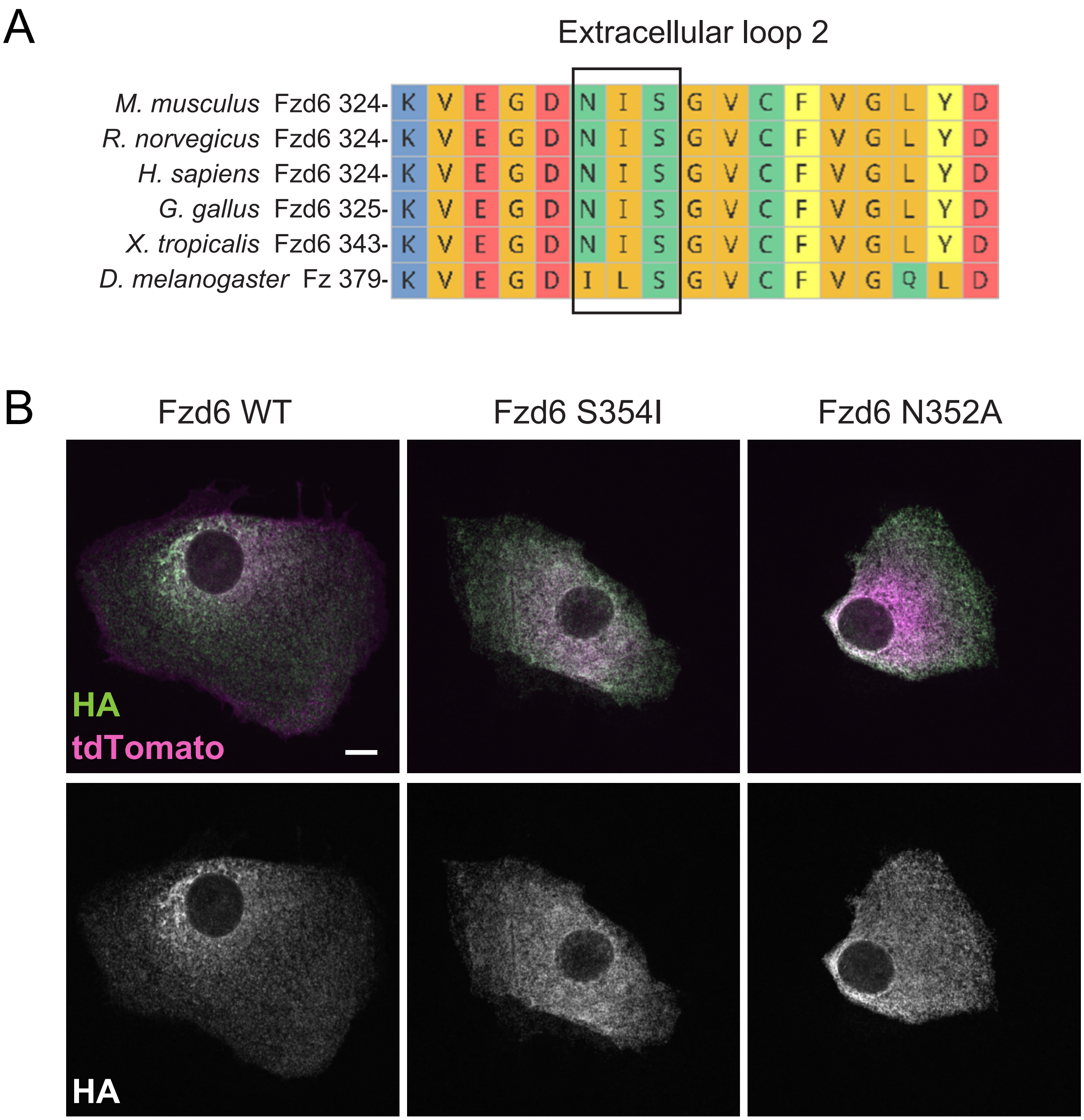


Figure S4. **The N-X-S consensus sequence is required for Fzd6 membrane localization.**

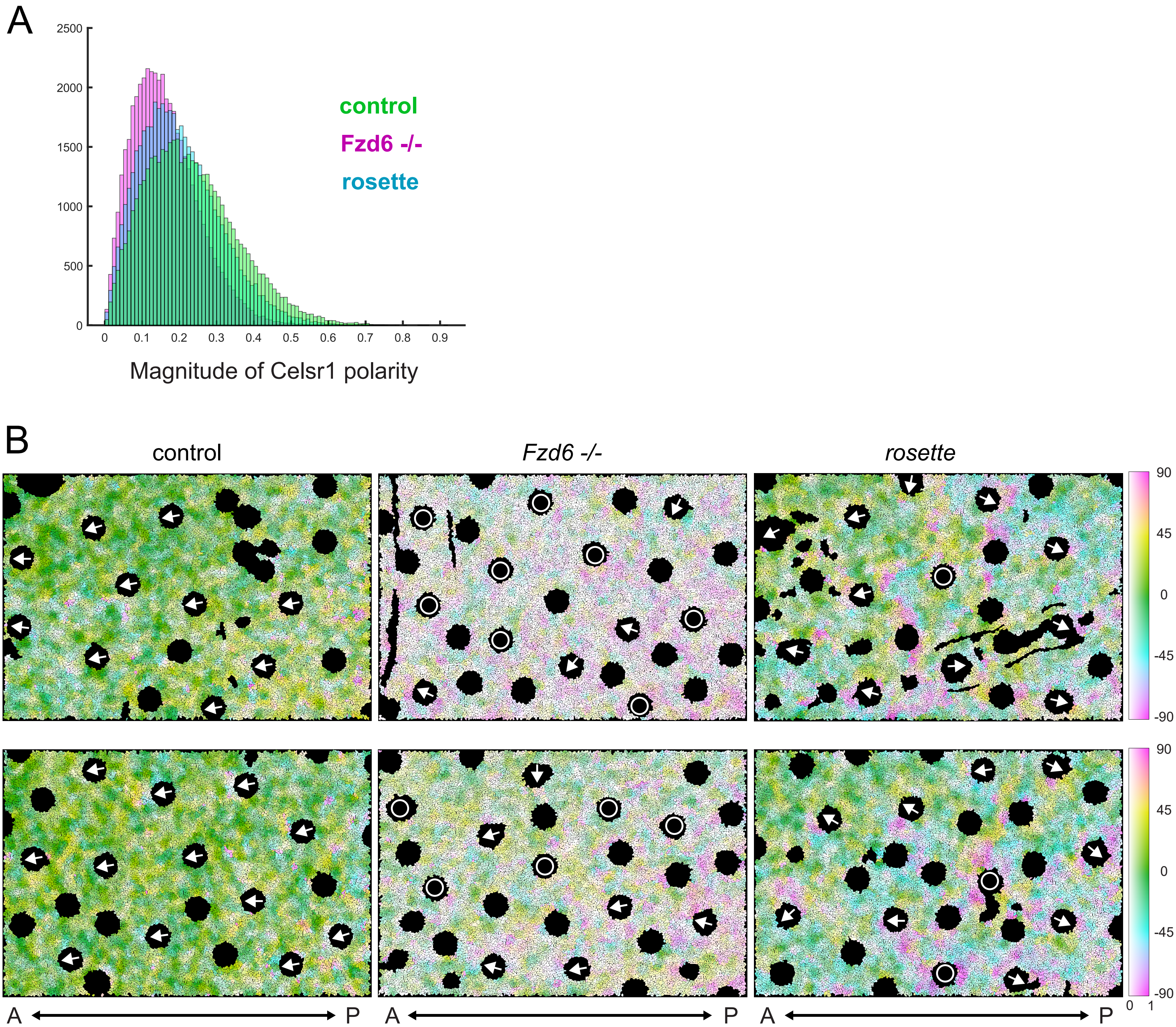


Figure S5. **The axis of PCP asymmetry rotates in the *rosette* epidermis.**

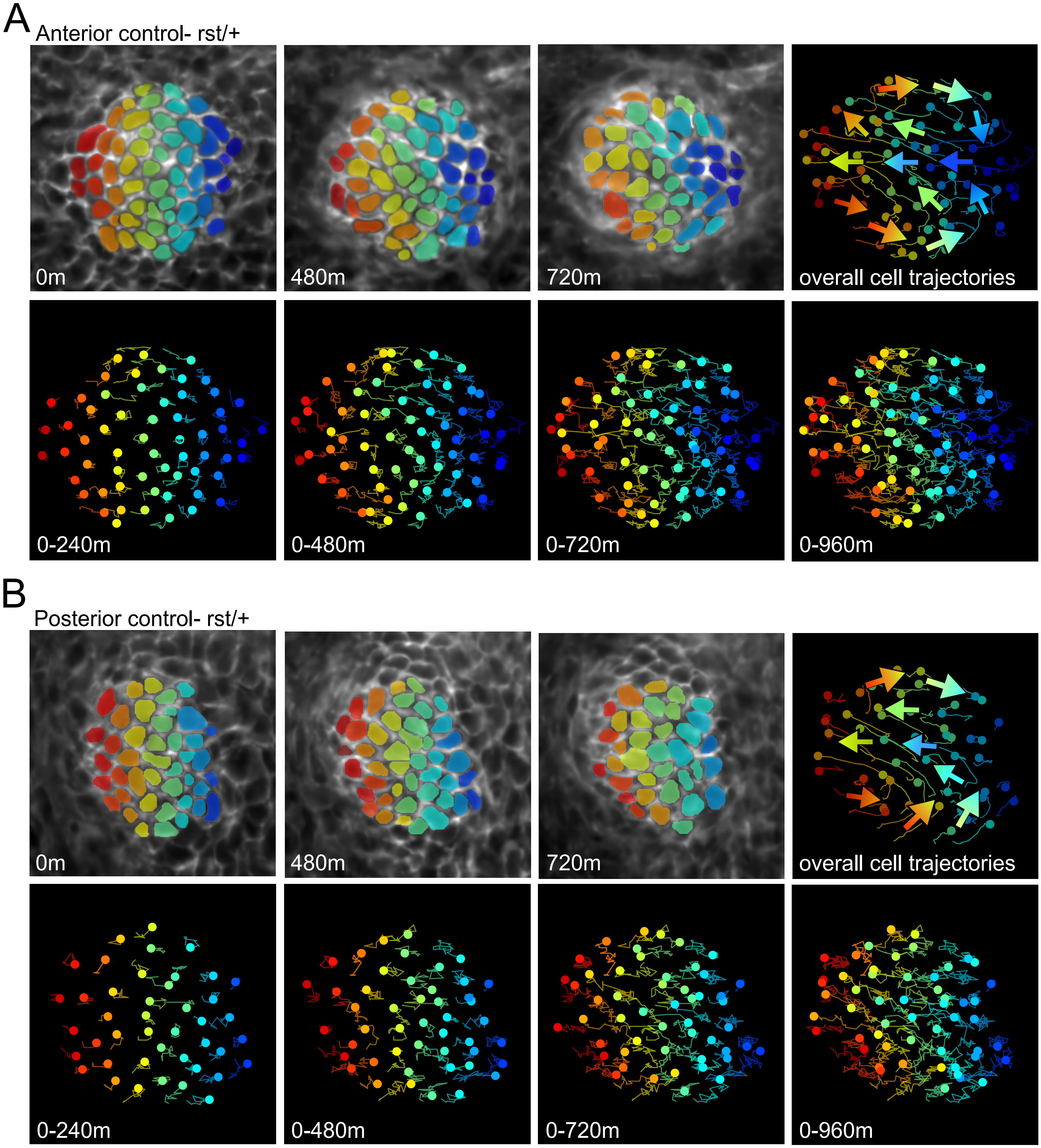


Figure S6. **PCP-directed collective cell movements occur normally in rst/+ hair placodes.**
