## Supplemental Table 1 for "Region-specific reversal of epidermal planar polarity in the fancy *rosette* mouse"

| Plate ID | Mouse ID | Gender | Breed | Family-cousin (A) or sibling (A+rest) | Family-Parental | % C57BL/6 | Phenotype | Duplicate ID |
|---|---|---|---|---|---|---|---|---|
| A01 | **11610-TGP1LLLRR Brn** | m | fancy, C57BL/6 | A | | 71.875 | 1 | |
| A02 | **11610-TGP2LLLRR** | m | fancy, C57BL/6 | A | | 71.875 | 1 | |
| A03 | **11610-TGP3LLLR** | m | fancy, C57BL/6 | A | | 71.875 | 1 | |
| A04 | **11610-TGP4LLLL** | m | fancy, C57BL/6 | A | | 71.875 | 1 | |
| A05 | **11610-TGP5LLRRRR** | f | fancy, C57BL/6 | A | I | 71.875 | 1 | |
| A06 | **11610-TGP6RRRLLL** | f | fancy, C57BL/6 | A | H | 71.875 | 1 | |
| A07 | **11610-TGP7LLLLRRR** | f | fancy, C57BL/6 | A | J | 71.875 | 1 | |
| A08 | **11610-TGP8RRRRL** | f | fancy, C57BL/6 | A | | 71.875 | 1 | |
| A09 | **11610-TGP9RRLLL** | f | fancy, C57BL/6 | A | | 71.875 | 1 | |
| A10 | **11610-TGP10RRRLLL** | f | fancy, C57BL/6 | A | G | 71.875 | 1 | |
| A11 | **11611-TGP1LLLRR brn** | m | fancy, C57BL/6 | A | | 71.875 | 1 | |
| A12 | **11611-TGP2LLLRR** | m | fancy, C57BL/6 | A | | 71.875 | 1 | |
| B01 | **11611-TGP3RRR** | m | fancy, C57BL/6 | A | | 71.875 | 1 | |
| B02 | **11611-TGP4RRL** | m | fancy, C57BL/6 | A | | 71.875 | 1 | |
| B03 | **11611-TGP5LLLRR brn** | f | fancy, C57BL/6 | A | F | 71.875 | 1 | |
| B04 | **11611-TGP6RRRLL** | f | fancy, C57BL/6 | A | | 71.875 | 1 | |
| B05 | **11760-TGP1LLLRRR** | m | fancy, C57BL/6 | A | J | 71.875 | 1 | |
| B06 | **11760-TGP2LLLRR** | m | fancy, C57BL/6 | A | I | 71.875 | 1 | |
| B07 | **11760-TGP3LLLRRR** | f | fancy, C57BL/6 | A | E | 71.875 | 1 | |
| B08 | **11760-TGP4RRRLL** | f | fancy, C57BL/6 | A | | 71.875 | 1 | |
| B09 | **11761-TGP1** | unknown | fancy, C57BL/6 | A | | 71.875 | 1 | |
| B10 | **11761-TGP2** | unknown | fancy, C57BL/6 | A | | 71.875 | 1 | |
| B11 | **11761-TGP3** | unknown | fancy, C57BL/6 | A | | 71.875 | 1 | |
| B12 | **11761-TGP4** | unknown | fancy, C57BL/6 | A | | 71.875 | 1 | |
| C01 | **11761-TGP5** | unknown | fancy, C57BL/6 | A | | 71.875 | 1 | |
| C02 | **11761-TGP6** | unknown | fancy, C57BL/6 | A | | 71.875 | 1 | |
| C03 | **11762-TGP1LLLBrn** | m | fancy, C57BL/6 | A | | 71.875 | 1 | |
| C04 | **11762-TGP2RRL** | m | fancy, C57BL/6 | A | | 71.875 | 1 | |
| C05 | **11762-TGP3LLLLRRRR** | m | fancy, C57BL/6 | A | | 71.875 | 1 | |
| C06 | **11762-TGP4LLLRRR** | f | fancy, C57BL/6 | A | | 71.875 | 1 | |
| C07 | **11610-TGN11LLLL** | m | fancy, C57BL/6 | A | | 71.875 | 0 | |
| C08 | **11610-TGN12LLRR** | m | fancy, C57BL/6 | A | | 71.875 | 0 | |
| C09 | **11610-TGN13RRRLL** | m | fancy, C57BL/6 | A | | 71.875 | 0 | |
| C10 | **11610-TGN14LLLRR** | m | fancy, C57BL/6 | A | | 71.875 | 0 | |
| C11 | **11610-TGN15LLLR** | f | fancy, C57BL/6 | A | | 71.875 | 0 | |
| C12 | **11610-TGN16LLRR** | f | fancy, C57BL/6 | A | | 71.875 | 0 | |
| D01 | **11610-TGN17LLLRR** | f | fancy, C57BL/6 | A | | 71.875 | 0 | |
| D02 | **11610-TGN18RRRLL** | f | fancy, C57BL/6 | A | | 71.875 | 0 | |
| D03 | **11611-TGN7LLLLR** | m | fancy, C57BL/6 | A | | 71.875 | 0 | |
| D04 | **11611-TGN8RRRLL** | m | fancy, C57BL/6 | A | | 71.875 | 0 | |
| D05 | **11611-TGN9LLLRR** | m | fancy, C57BL/6 | A | | 71.875 | 0 | |
| D06 | **11611-TGN10LLLRR** | f | fancy, C57BL/6 | A | | 71.875 | 0 | |
| D07 | **11611-TGN11RRRRL** | f | fancy, C57BL/6 | A | | 71.875 | 0 | |
| D08 | **11611-TGN12LLLLR** | f | fancy, C57BL/6 | A | | 71.875 | 0 | |
| D09 | **11611-TGN13LLLRR** | f | fancy, C57BL/6 | A | | 71.875 | 0 | |
| D10 | **11611-TGN14RRRRL** | f | fancy, C57BL/6 | A | | 71.875 | 0 | |
| D11 | **11760-TGN6LLLLRRR** | m | fancy, C57BL/6 | A | | 71.875 | 0 | |
| D12 | **11760-TGN7RRRRLLL** | m | fancy, C57BL/6 | A | | 71.875 | 0 | |
| E01 | **11760-TGN8RRL** | m | fancy, C57BL/6 | A | | 71.875 | 0 | |
| E02 | **11760-TGN9RRRRLLL** | f | fancy, C57BL/6 | A | | 71.875 | 0 | |
| E03 | **11760-TGN10LLLLRRR** | f | fancy, C57BL/6 | A | | 71.875 | 0 | |
| E04 | **11760-TGN11RRRRLLL** | f | fancy, C57BL/6 | A | | 71.875 | 0 | |
| E05 | **11762-TGN1RRR** | f | fancy, C57BL/6 | A | | 71.875 | 0 | |
| E06 | **11762-TGN2RRLBrn** | f | fancy, C57BL/6 | A | | 71.875 | 0 | |
| E07 | **11762-TGN3LLLRRRR** | f | fancy, C57BL/6 | A | | 71.875 | 0 | |
| E08 | **11762-TGN4LLLLRRR** | f | fancy, C57BL/6 | A | | 71.875 | 0 | |
| E09 | **original stud** | m | fancy | B | FOUNDER | 0 | 1 | H06 |
| E10 | **1031-1blkF3RRRRLLL** | m | fancy, C57BL/6 | | A,C,D,H, K, L | 62.5 | 1 | H07 |
| E11 | **105801F4RRRRLL** | f | fancy, C57BL/6 | C | A | 81.25 | 0 | H08 |
| E12 | **105802F4LLLLRR** | f | fancy, C57BL/6 | C | A | 81.25 | 0 | H09 |
| F01 | **10370-1RRLLLL** | f | fancy, C57BL/6 | D | | 56.25 | 1 | |
| F02 | **10370-2LLLRRR** | f | fancy, C57BL/6 | D | | 56.25 | 1 | |
| F03 | **10370-3LLRRRR** | m | fancy, C57BL/6 | D | E | 56.25 | 1 | |
| F04 | **10370-5LLLRRRR** | m | fancy, C57BL/6 | D | F,G | 56.25 | 1 | |
| F05 | **12070-1LLLR** | m | fancy, C57BL/6 | E | | 64.0625 | 1 | |
| F06 | **12070-2LLLLRRR** | f | fancy, C57BL/6 | E | | 64.0625 | 1 | |
| F07 | **12070-3LLLLRRRR** | f | fancy, C57BL/6 | E | | 64.0625 | 1 | |
| F08 | **12070-4RRR** | f | fancy, C57BL/6 | E | | 64.0625 | 1 | |
| F09 | **12070-5RRRRLLL** | f | fancy, C57BL/6 | E | | 64.0625 | 1 | |
| F10 | **12070-6LLRRBrn** | f | fancy, C57BL/6 | E | | 64.0625 | 1 | |
| F11 | **12090-1LLRRR** | m | fancy, C57BL/6 | F | | 64.0625 | 1 | |
| F12 | **12100-1RRL** | m | fancy, C57BL/6 | G | | 64.0625 | 1 | |
| G01 | **12100-2RRRRLLL** | m | fancy, C57BL/6 | G | | 64.0625 | 1 | |
| G02 | **12100-3RRLLLL** | f | fancy, C57BL/6 | G | | 64.0625 | 1 | |
| G03 | **12100-4RRRLL** | f | fancy, C57BL/6 | G | | 64.0625 | 1 | |
| G04 | **12100-5LLLRR** | f | fancy, C57BL/6 | G | | 64.0625 | 1 | |
| G05 | **12120-1RRRRLLLL** | m | fancy, C57BL/6 | H | | 67.1875 | 1 | |
| G06 | **12120-2RRR** | f | fancy, C57BL/6 | H | | 67.1875 | 1 | |
| G07 | **12120-3RRLBrn** | f | fancy, C57BL/6 | H | | 67.1875 | 1 | |
| G08 | **12120-4LLLLRRR** | f | fancy, C57BL/6 | H | | 67.1875 | 1 | |
| G09 | **12120-5LLLR** | f | fancy, C57BL/6 | H | | 67.1875 | 1 | |
| G10 | **12130-1RRLLL** | m | fancy, C57BL/6 | I | | 71.875 | 1 | |
| G11 | **12130-2RRRLL** | m | fancy, C57BL/6 | I | | 71.875 | 1 | |
| G12 | **12150-1LLLLRRR** | f | fancy, C57BL/6 | J | | 71.875 | 1 | |
| H01 | **12151-1RRRLL** | m | fancy, C57BL/6 | J | | 71.875 | 1 | |
| H02 | **10390-1LLLLR** | f | fancy, C57BL/6 | K | | 56.25 | 1 | |
| H03 | **10720-1LRRRR** | m | fancy, C57BL/6 | L | | 56.25 | 1 | |
| H04 | **10720-2LLLLR** | m | fancy, C57BL/6 | L | | 56.25 | 1 | |
| H05 | **original brother** | m | fancy | B | | 0 | 1 | |
| H06 | **original stud** | m | fancy | B | FOUNDER | 0 | 1 | E09 |
| H07 | **1031-1blkF3RRRRLLL** | m | fancy, C57BL/6 | | A,C,D,H, K, L | 62.5 | 1 | E10 |
| H08 | **105801F4RRRRLL** | f | fancy, C57BL/6 | C | A | 81.25 | 0 | E11 |
| H09 | **105802F4LLLLRR** | f | fancy, C57BL/6 | C | A | 81.25 | 0 | E12 |
| H10 | **B6 female** | f | C57BL/6 | | | 100 | 0 | H11 |
| H11 | **B6 female** | f | C57BL/6 | | | 100 | 0 | H10 |
| H12 | **B6 male** | m | C57BL/6 | | | 100 | 0 | |
